# Discovering Polyphosphate and Polyhydroxyalkanoate-Accumulating Organisms Across Ecosystems: Phenotype-targeted Genotyping via FACS-sequencing

**DOI:** 10.1101/2025.03.24.644850

**Authors:** Yuan Yan, Mathew Baldwin, Jangho Lee, Zijian Wang, Guangyu Li, Il Han, Peisheng He, Varun Shrinivasan, Liam Wickes-Do, Michael A. Gore, April Z. Gu

## Abstract

Intracellular biopolymers serve versatile functions that allow microbes to adapt to fluctuating environmental conditions. The metabolic interdependence of dual intracellular polymers, namely polyphosphate (polyP) and polyhydroxyalkanoates (PHA), is a defining feature of functionally polyphosphate-accumulating organisms (PAOs), the key agents enabling enhanced biological phosphorus removal (EBPR) for wastewater treatment. However, beyond EBPR systems, the presence and identities of PAOs that possess both polyP and PHA in natural environments such as soil have never been examined due to a lack of available detection tools, despite their potential roles in carbon and phosphorus cycling. This study presents a novel phenotype-targeted approach integrating triple-stained fluorescence-activated cell sorting (FACS) with 16S rRNA gene amplicon sequencing (termed TriFlow-Seq) to simultaneously detect, quantify, and phylogenetically characterize PAOs accumulating both polyP and PHA (referred to as PHA- PAOs). TriFlow-Seq was validated using polymer staining image analysis and Single-Cell Raman micro-spectroscopy. Application to EBPR systems successfully enriched known PHA- PAOs, including *Candidatus* Accumulibacter, *Tetrasphaera*, *Dechloromonas*, *Pseudomonas.* It also revealed novel candidate PHA-PAOs, particularly within the Rhodobacteraceae family. When applied to soil samples, TriFlow-Seq led to the first discovery of diverse PHA-PAOs dominated by *Pseudomonas*, *Halomonas*, and *Nannocystis* in maize rhizosphere soils. These predominant genera are known rhizosphere inhabitants of essential crops with key plant growth-promoting functions including phosphate solubilization, biofilm formation, and phytohormone production, yet simultaneous polyP and PHA accumulation has not been directly reported in the maize rhizosphere. Our findings revealed unexpectedly high PHA-PAO prevalence and distinct phylogenetic patterns associated with different maize genotypes, suggesting a potentially overlooked role for dual polymer storage in microbial rhizosphere dynamics and function. This study establishes a pioneering approach to investigate dual polyP and PHA-containing PAO identities and their important roles in rhizosphere structure and plant health.

**Synopsis:** Novel method reveals dual-polymer bacteria in activated sludge and crop rhizospheres, suggesting new candidates supporting wastewater treatment and agricultural sustainability.

## 1 Introduction

A diverse range of microbes contains intracellular biopolymers, including polyphosphate (polyP), polyhydroxyalkanoates (PHA), glycogen, waxy esters, triacylglycerides (TAGs), trehalose, and cyanophycin.^1^ These biopolymers serve versatile functions such as energy storage, nutrient cycling, ion balance, redox homeostasis, and stress resistance, allowing microbes to adapt to fluctuating environmental conditions.^1–3^ Among them, polyP-accumulating organisms (PAOs) are key agents in achieving sustainable phosphorus (P) removal and recovery in enhanced biological phosphorus removal (EBPR) systems.^4,5^ PAOs exhibit a unique metabolism: under anaerobic conditions, they utilize stored polyP as an energy source, releasing phosphate and concurrently accumulating PHA. Then, under subsequent oxic or anoxic conditions, they uptake phosphate and synthesize new polyP while using stored PHA to fuel growth.^6^ This unique metabolism provides competitive advantages for PAOs under fluctuating anaerobic/oxic conditions.^7–9^ Therefore, the metabolic interdependence of dual intracellular polymers (polyP and PHA) is a defining feature of functionally verified PAOs in EBPR systems, such as *Candidatus* Accumulibacter,^6^ *Pseudomonas*,^10^ and *Dechloromonas;*^7^ although some, such as *Tetrasphaera,* have distinct sub-groups with varying capacities to accumulate PHA.^11–14^

Beyond EBPR engineering systems, there is increasing interest in exploring the functional roles of PAOs in natural environments such as soil. Emerging evidence points to their ubiquitous presence and versatile functions, where they serve as important facilitators of phosphorus (P) availability, carbon (C) cycling, biofilm formation, and contribute to both plant and microbial community stress resilience.^15–18^ In the natural environments, PAOs are often broadly identified as polyP-containing organisms, with little knowledge of the potential co-occurrence and co-metabolisms with C biopolymer storage. To explore this, we performed a preliminary quantification of polyP-containing organisms (PCOs) in agricultural soil with single-cell Raman spectroscopy analysis, which indicated that up to 73.7% of PCOs also contain C storage biopolymers, with a significant Pearson’s correlation between PCOs and C-accumulating PCOs (*r*=0.8, *p*-value <0.001) (Figure S1). While exploratory, this unexpected observation of a high occurrence of PAOs simultaneously possessing both polyP and PHA suggests that dual storage may be an ecologically relevant functional trait of PAOs in soil. It highlights complex intra- and interdependent metabolic pathways that remain poorly understood. As the most common C storage biopolymer, PHA (∼1-4 μg C/g soil) responds positively to C availability, but is suppressed by nitrogen (N) availability, suggesting its role in stoichiometric balancing and stress response as an alternative carbon allocation strategy.^19,20^ The co-storage of C biopolymers and polyP likely enhances microbial adaptability to nutrient fluctuations and contributes to broader nutrient cycling and ecosystem functioning. This dual-storage strategy distinguishes specialized PAOs that exhibit resource-dependent surplus storage from organisms that accumulate relatively low level of polyP primarily for stress responses or cation sequestration.^20–22^ Fungal PAOs have been more extensively studied in soil,^23^ whereas little is known about bacterial PAOs in managed environments. The inability of polyP-synthesizing fungal organisms, such as arbuscular mycorrhizal fungi (AMF), to produce PHA provides a clear demarcation between bacterial and fungal PAOs. Although there have been emerging yet limited investigations of the presence and roles of PAOs in nature environments, the presence and identities of bacterial PHA accumulating PAOs (PHA-PAOs) in natural environments have not yet been elucidated. Particularly, we are interested in identifying bacterial microorganisms exhibiting surplus storage, defined by their specific ability to accumulate substantial polyP and PHA beyond immediate biological maintenance, distinguishing them from microbes that maintain constitutively low levels of these biopolymers for basic cellular functions (e.g., *Escherichia coli* or *Bacillus subtilis*).^24,25^

Despite the ecological, biogeochemical, and engineering significance of PHA-PAOs, identifying functionally active and phylogenetically novel PAOs exhibiting both intracellular polyp and PHA storage in complex environmental matrices remains challenging.^15,26–28^ Methods for detecting PAOs that co-accumulate polyP and PHA in natural systems have yet to be reported. Conventional approaches rely heavily on gene-centric profiling such as polyP-metabolism biomarkers (i.e., *ppk*) or metagenomics to identify and quantify previously known PAOs across aquatic and terrestrial systems,^29,30^ yet these methods have limitations and cannot reflect actual phenotypic traits such as active polyP and/or PHA accumulation and co-accumulation. ^25–27^ Discovery of novel PAOs is often hindered by *ppk* gene-based detection which relies on prior genetic information of the target PAOs. Additionally, these methods have low specificity for detecting PAOs with unique polyP accumulation abilities, as the polyP-synthesizing gene *ppk* is ubiquitous across microorganisms.^31,32^ Moreover, polyP metabolism is not solely regulated by the *ppk* gene, as alternative proteins, such as actin-like Arp proteins, have been proposed to exhibit PPK-like functions in bacteria.^33^ A few recent studies have employed fluorescence-activated cell sorting (FACS) to phenotypically track the polyP accumulation traits and resolve the phylogeny and taxonomy of PAOs from EBPR systems.^34,35^ However, these studies used stains only for polyP, without detecting the co-metabolism polymers like PHA.^34^ In addition, FACS-based PAO detection in soil samples has never been reported.

In this study, we propose a refined definition of functional PAOs in natural environments as organisms that co-accumulate both polyP and intracellular carbon, predominantly PHA, to surplus levels, where this co-metabolism holds important ecological roles as discussed above, and here refer them as PHA-PAOs. We developed TriFlow, a workflow integrating triple-color fluorescence staining with FACS to enable the simultaneous detection and isolation of functionally active PAOs that co-accumulate both polyP and PHA. When coupled with 16S ribosomal RNA (16S rRNA) amplicon sequencing for phylogenetic characterization, the workflow becomes TriFlow-Seq. This extended method links phenotypic traits to taxonomic identities. We validated TriFlow by comparing the relative abundances of PHA-PAOs detected using TriFlow and single cell Raman spectroscopy (SCRS), an established method for the simultaneous detection of intracellular polyP and PHA in PAOs within EBPR systems.^27,36–39^ We further demonstrated the applicability of TriFlow-Seq across diverse environmental samples, including activated sludge from four full-scale and one pilot-scale EBPR facilities known to contain PHA-PAOs, as well as rhizosphere soil from experimental maize fields in New York. This new method enables us to detect and identify previously unknown bacterial PHA-PAOs and provide targets for further targeted investigations of novel metabolisms and functional roles of these unique and prevalent functional groups of organisms across diverse environmental systems.

## 2 Material and Methods

### 2.1 Sample collection

#### 2.1.1 Activated sludge samples from EBPR systems

Biomass samples from the aerobic zone were collected from three full-scale and one pilot-scale plant, all of which effectively remove P and contain active PHA-PAOs. Activated sludge samples from full-scale EBPR systems were stored at 4°C, transported overnight to the laboratory, and activated with a phosphorus release/uptake test (Text S1, Figure S2), while sludge samples from the pilot-scale S2EBPR were directly fixed onsite as described in Section 2.3.

#### 2.1.2 Soil samples

In 2023, two sets of maize genotypes—inbred and hybrid lines (Table S1)—were planted at Cornell University’s Musgrave Research Farm in Aurora, NY. The set of inbred lines included five of the 26 parents of the U.S. nested association mapping population,^40^ as well as both parents of the intermated B73 × Mo17 population,^41^ with B73 serving as a parent in both populations. Collectively, the inbred set included popcorn, sweet corn, temperate, and tropical maize inbred lines. The hybrid set consisted of 22 temperate-adapted hybrids that represented the high-intensity phenotyping site plots provided by the Genomes to Fields (G2F) Initiative (https://www.genomes2fields.org/home/); however, only 10 of the 22 hybrids were sampled in this experiment.

The inbred set was planted on May 18 in a field (42.723288, −76.657356) characterized by a Lima silt loam (0–3% slope), while the hybrid set was planted on May 12 in a field (42.733297, −76.653955) characterized by both Honeoye silt loam (3–8% slopes) and Kendaia & Lyons silt loams (0–3% slope). Both genotype sets were planted in a randomized complete block design with two replicates, with experimental units consisting of two-row plots of varying lengths at each field location.

Plot dimensions and planting rates are described in Table S1. Rhizosphere samples were collected at tasseling (VT) in accordance with Hanway et al.^36^ Briefly, within each two-row plot, one plant per row was selected at random and uprooted with bulk soil gently shaken off; the remaining soil adhering to the roots was collected as rhizosphere soil. However, due to greater planting density in hybrid plots, three plants per two-row plot (split between the two rows) were sampled. Samples were pooled by plot (n=2 per genotype), stored, and transported at 4 C for sample pre-processing within 24 hours.

### 2.2 Microbiota isolation

#### 2.2.1 Activated sludge

Sludge samples were collected from the middle of the aerobic phase and centrifuged to remove the supernatant. The remaining pellets were washed three times with 1 × PBS to remove any excess cations. All centrifuge steps for activated sludge samples were carried out at 3200 × g for 10 minutes due to the easy settleability unless otherwise specified.

#### 2.2.2 Rhizosphere microbiota

Extensive sample pre-treatment and a targeted sequential gating strategy were developed to separate, disaggregate, and isolate PHA-PAOs at the single-cell scale from the soil matrix. The key steps for isolating PHA-PAOs and ensuring single-cell recovery included: 1) detergent-based cell detachment and subsequent cell separation based on sedimentation rate and filtering, 2) disaggregation via chemical and enzymatic washing to destabilize soil microbial aggregates, 3) identification and sorting during gating (Text S2) based on particle size using forward scatter height and area, and 4) triple-fluorescence indicating simultaneous accumulation of DNA, PHA, and polyP.

##### 2.2.2.1 Physical-chemical cell detachment

To isolate the microbial fraction from the soil matrix, 1 gram of fresh soil was suspended in 20 mL of a detachment solution containing 3 mM sodium pyrophosphate, 0.5% Tween 20, and 0.35% polyvinylpyrrolidone in 1 × phosphate-buffered saline (PBS) (0.4 M Na HPO /NaH PO, 150 mM NaCl, pH 7.2).^43^ The soil suspension was agitated for 30 minutes at room temperature in accordance with Deng et al. 2019,^44^ at 200 rpm using an orbital shaker to facilitate detergent based cell detachment (Figure S3, step 2).

##### 2.2.2.2 Cell separation

Following cell detachment on orbital shaker, to enhance cell recovery, the resulting soil slurry was vortexed for 10 seconds and left to settle for one minute to sediment larger soil particles (Figure S3, step 3). Due to previously reported wide range of sedimentation rates of soils, known to vary based on soil texture, physicochemical properties, and particle size, settling time should be adjusted for individual samples.^45^ For these soil samples, the sedimentation time was optimized as 1 minute based on preliminary time series analysis used to monitor the sedimentation process (Figure S4). Following sedimentation, cell separation from the soil matrix was further enhanced by decanting through a 40 μm cell strainer (Corning®, USA, Cat. No. 431750). Here we acknowledge we are unable to directly isolate and ensure cell purity in the decanted supernatant following filtration due to the small size of many particles within soils; we further address cell purity during sorting in our gating strategy in Text S2. Decanted and filtered cell suspension was then pelleted via centrifugation at 6000 × g for 10 minutes with remaining supernatant decanted (Figure S3, step 3). All centrifuge steps for rhizosphere samples were carried out at 6000 × g for 10 minutes unless otherwise specified.

##### 2.2.2.3 Cell Cleaning

To enhance microbial dispersion for downstream fluorescent staining and FACS, the detached, filtered, and pelleted soil community was resuspended and washed once with 5 mL of 4 mM ethylenediaminetetraacetic acid (EDTA) in 1 × PBS (pH ∼7.2) to remove excess cations which facilitate aggregation of microbial cells in electrostatic interactions with: microbial cell walls, extracellular polymeric substances (EPS).^46^ This was followed by two additional washings with 5mL of 1 × PBS to remove any residual EDTA and solubilized cations (Figure S3, step 4).

### 2.3 Cell fixation

After microbial isolation in step 2.2, for both soil communities and sludge, the remaining sample pellets were resuspended in 5 mL of fixation buffer, vortexed for 10 s, and stored at 4°C for 24 hours. The fixation buffer was prepared according to previously described protocols, containing 5 mM nickel chloride (NiCl_2_), 5 mM barium chloride (BaCl_2_), 10% sodium azide (NaN_3_) in 4-(2- hydroxyethyl)-1-piperazineethanesulfonic acid (HEPES) buffer (25 mM, pH 7.2) (Figure S3, step 5).^35,47^ Previous efforts have noted no significant difference in fixed microbial community patterns within 9 days.^47^

### 2.4 Triple-staining

Following 24 hours of fixation, both sludge and soil samples were pelleted by centrifugation and washed three times with 1× PBS to remove residual fixative.

Fixed and washed activated sludge samples need an additional homogenization step: sludge was mechanically disrupted with a 26-gauge needle and 1 mL syringe by repeatedly drawing the sludge into the syringe and out vigorously for 5 mins. Compared to soil, activated sludge has a simpler composition with fewer inorganic particles, making it more amenable to single-cell separation by shear. This approach, adapted from established SCRS protocols, effectively isolates individual cells for analysis.^36,37^ After homogenization for 5 min, sludge samples were washed an additional three times with 1 × PBS.

After washing steps, both soil and sludge samples were resuspended to 5 mL with 1 × PBS and treated with 1.6 mL solution A (0.11 M citric acid/4.1 Mm Tween 20 in distilled water) for 20 min to permeabilize the cells, then washed and resuspended in 5 mL of 1 × PBS, and filtered through a 40 μm cell strainer (Figure S3, step 6).

Deoxyribonucleic acid (DNA), polyP, and PHA of the cells were stained with SYBR Green I (SG) (Ex 488 nm / Em 515 nm) (Invitrogen, USA), tetracycline hydrochloride (TC) (Ex 390 nm / Em 560 nm) (Sigma Chemical Co., USA), and nile red (NR) (Ex 532 nm / Em 610 nm) (Sigma Chemical Co., USA), respectively–each validated in previous studies.^35,48,49^ TC was first applied at 2 mg/mL in Milli-Q water (final concentration 0.225 mM in cell suspensions) and incubated for 10 minutes.^35^ NR stock (10 mg/mL in DMSO) was added to 1 mL of cell suspension at a final concentration of 0.05 mg/mL and incubated for 10 minutes.^50^ Lastly, 100 × SG was added to the cells and incubated for 4 to 12 hours (Figure S3, step 7).^44^ All incubation steps were performed at room temperature in the dark.

### 2.5 Soil disaggregation

Significant challenges were faced when applying the FACS step of the TriFlow-Seq workflow to soil samples, which differ markedly from activated sludge in their physicochemical properties and tendency to reaggregate. Traditional soil pretreatments using detergents or sonication for quantification have proven inadequate for sorting low-abundance populations,^51,52^ especially when high cell numbers are required for quality genomic reads. Key challenges include reaggregation during extended suspension, which reduces sorting efficiency and increases false positives,^53^ and soil heterogeneity, which affects reaggregation rates due to variations in cation exchange capacity, plant and microbial protein activity, soil organic matter content, and biofilm formation.^54^ This re-aggregation can reduce sorting efficiency and increase the risk of nozzle clogging, potentially contaminating ongoing cell sorting.^52^ To address this, a gentle enzymatic digestion using Trypsin-EDTA was applied after staining to degrade residual and persistent extracellular organic materials in microbial pellets, thereby further preventing re-aggregation during FACS.^55^ The protocol involved firstly pelleting the stained sample by centrifugation after incubation with SG and decanting the supernatant. The stained pellets were then resuspended in 3 mL of 1 × Trypsin-0.5 mM EDTA and incubated for 1 min at 37°C in a water bath, followed by trypsin deactivation using 1 × FBS in a 1:1 ratio. Prior to FACS, trypsin treated samples were diluted in 1 × PBS to achieve a proper threshold rate of 1500 events s^-1^ (Figure S3, step 8). Notably, this additional pretreatment step achieved sorting efficiencies ranging from 77-92% with an average of 85.5% (Table S2) across the tested soil series, representing a substantial improvement of approximately 50% higher compared to the 35.75% average sorting efficiency obtained using activated sludge sample pretreatment (n = 8 samples, Table S2).

### 2.6 FACS instrument settings

Cell sorting was carried out using the 100-micron nozzle on a Fusion Aria III cell sorter (BD, U.S.) equipped with a blue laser (488 nm excitation) with a 530 nm band-pass (BP) filter for SG, a violet laser (405 nm excitation) with a 525 BP filter for TC, and a yellow-green laser (561 nm excitation) with a 610 BP filter for NR. The blue laser was also used to analyze forward scatter (FSC) and side scatter (SSC), corresponding to cell size and cell granularity, respectively. Manual gating was performed as summarized in the supplementary material (Text S2, Figure S3). Bacterial cells were sorted in purity mode at a rate below 1,500 events/second. To obtain sufficient DNA for downstream analyses, up to 10 cells were sorted into sterile 1.5 mL microtubes and stored at −80°C for further analysis.^56^

### 2.7 FACS gating validation

A post-sort purity check was conducted to confirm single-cell isolation and targeted sorting of PHA-PAOs. A 100 μl aliquot of sorted sample was diluted in 1 × PBS and re-analyzed. Approximately 89% of events fell within the FSC-H/FSC-A gate, indicating successful recovery of single cells, while around 57% remained within the PHA-PAO gate—consistent with expectations due to photobleaching during laser exposure.

To ensure the reliability and accuracy of the proposed TriFlow-Seq method in identifying and quantifying single cell PHA-PAOs in complex microbial communities, we employed two independent validation methods. First, we verified PHA-PAO enrichment via fluorescent confocal microscopy. Approximately 4,000 sorted cells were collected to validate that all sorted cells were positive for SG, TC, and NR. Unsorted cells (as a control) and sorted cells were first filtered through a black polycarbonate filter (0.22 µm pore size) to concentrate the cells. The filters were placed onto glass slides with a drop of antifade reagent. Validation of the gating methods was carried out using confocal microscopy (Carl Zeiss) at 800X magnification with three excitation/filter sets: blue light (bandpass (BP) 450–490 nm excitation filter, dichroic beamsplitter (FT) at 500 nm, and longpass (LP) 515 nm emission filter), green light (BP 510– 560 nm, FT 565 nm, LP 590 nm), and ultraviolet (BP 340–380 nm, FT 400 nm, BP 435–485). Quantification of TC- and NR-stained cells relative to SG-stained cells was performed using Daime (version 1.2).^57^

### 2.8 DNA extraction and amplicon sequencing

Genomic DNA from sorted soil and sludge cells was extracted using the InstaGene Matrix (Bio-Rad Laboratories, USA). Genomic DNA from unsorted raw soil and sludge was extracted using the DNeasy PowerSoil Pro Kit (Qiagen, Valencia, CA, USA) on the QIAcube® (QIAGEN) platform. The V4 region of the 16S rRNA genes from both sorted and unsorted samples was amplified and sequenced using primers 515F and 806R at the University of Connecticut- Microbial Analysis, Resources, and Services (UCONN-MARS) facility. Details of the sequencing and analysis processes are listed in Text S3.

### 2.9 Classification of Detected Genera by Polyphosphate and PHA Accumulation Potential

For each detected genus, PAO-related functionality was assessed through literature review and genomic analysis. A color-coding system was applied: red stars indicated genera with experimentally confirmed anaerobic phosphorus release and aerobic phosphorus uptake; yellow stars denoted genera with documented polyP and PHA accumulation but unclear PAO activity; green stars represented genera with genetic potential for PHA-PAO metabolism based on functional gene presence in representative genomes from NCBI and BioCyc databases; blue stars were assigned when taxonomic resolution was limited to family or order level, with functional inference based on constituent genera. This color code system was applied in Figure 3 and 4, and Table S3 and S4. Detailed methodology is provided in Text S4.

**Figure 1.**
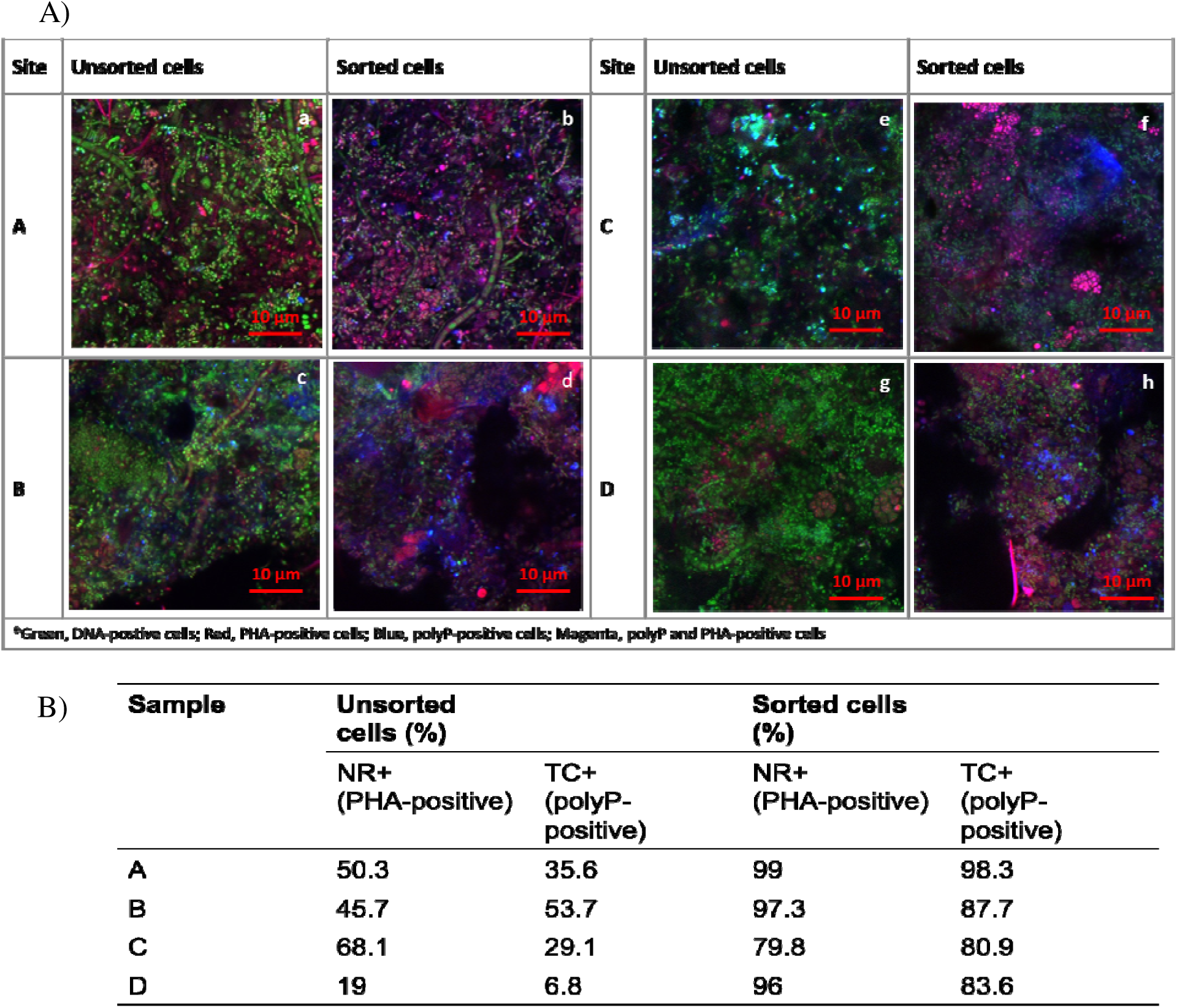
A) Confocal microscopy images comparing FACS sorted stain-positive cells to unsorted cells in activated sludge. Samples were taken from three EBPR systems (A, B, and C), and a S2EBPR system (D). B) Relative abundances of NR (PHA)-positive and TC (polyP)-positive cells in sorted samples versus original unsorted samples, as determined by semi-quantitative confocal microscopic image analysis.

**Figure 2.**
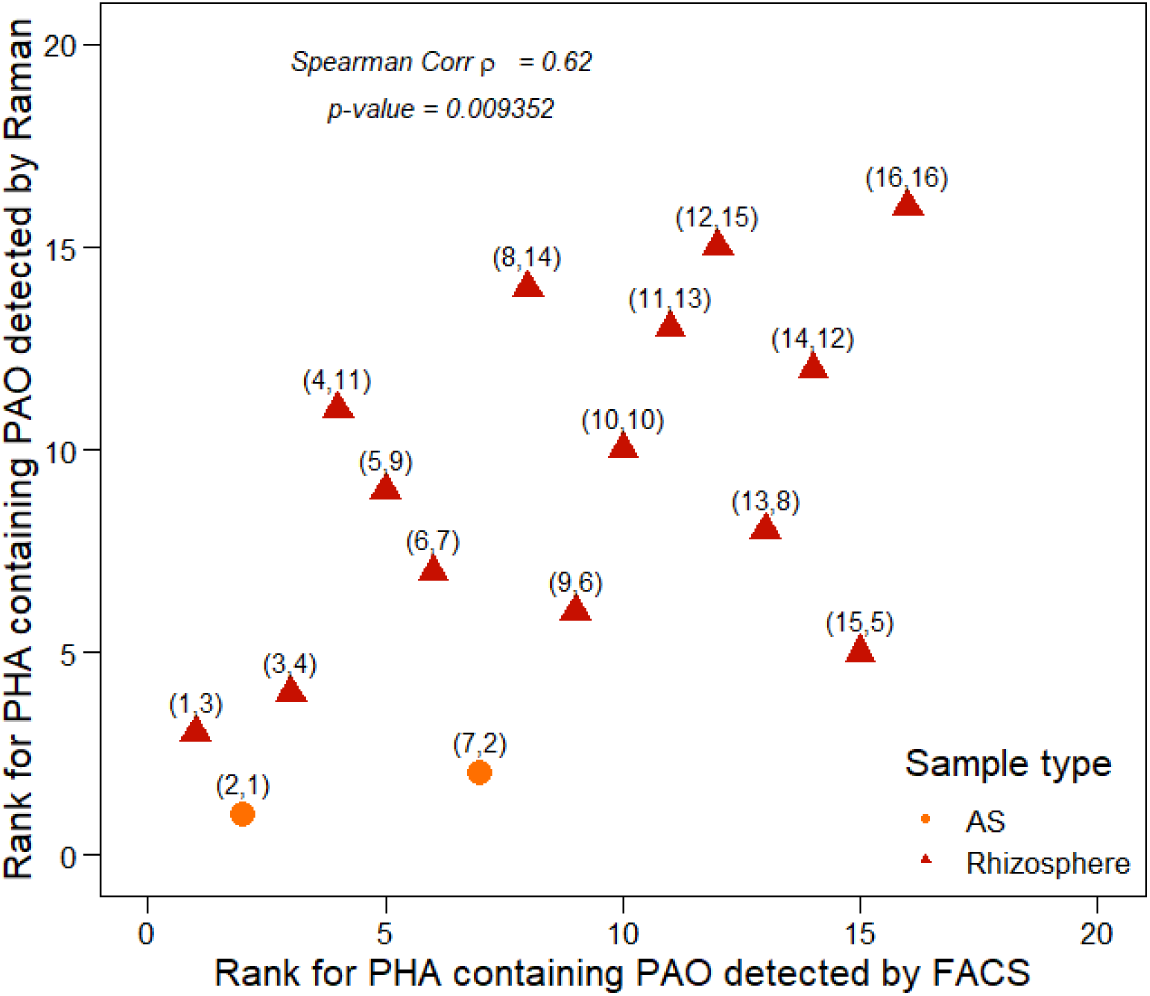
Spearman’s rank correlation coefficients of PHA-PAOs detected by FACS and SCRS. PHA-PAOs are defined as PHA-PAO cells containing both polyP and PHA. The strong correlation between the two independent techniques validates the effectiveness of TriFlow in detecting and quantifying PHA-PAOs in complex microbial communities. AS denotes activated sludge from EBPR and S2EBPR systems.

**Figure 3.**
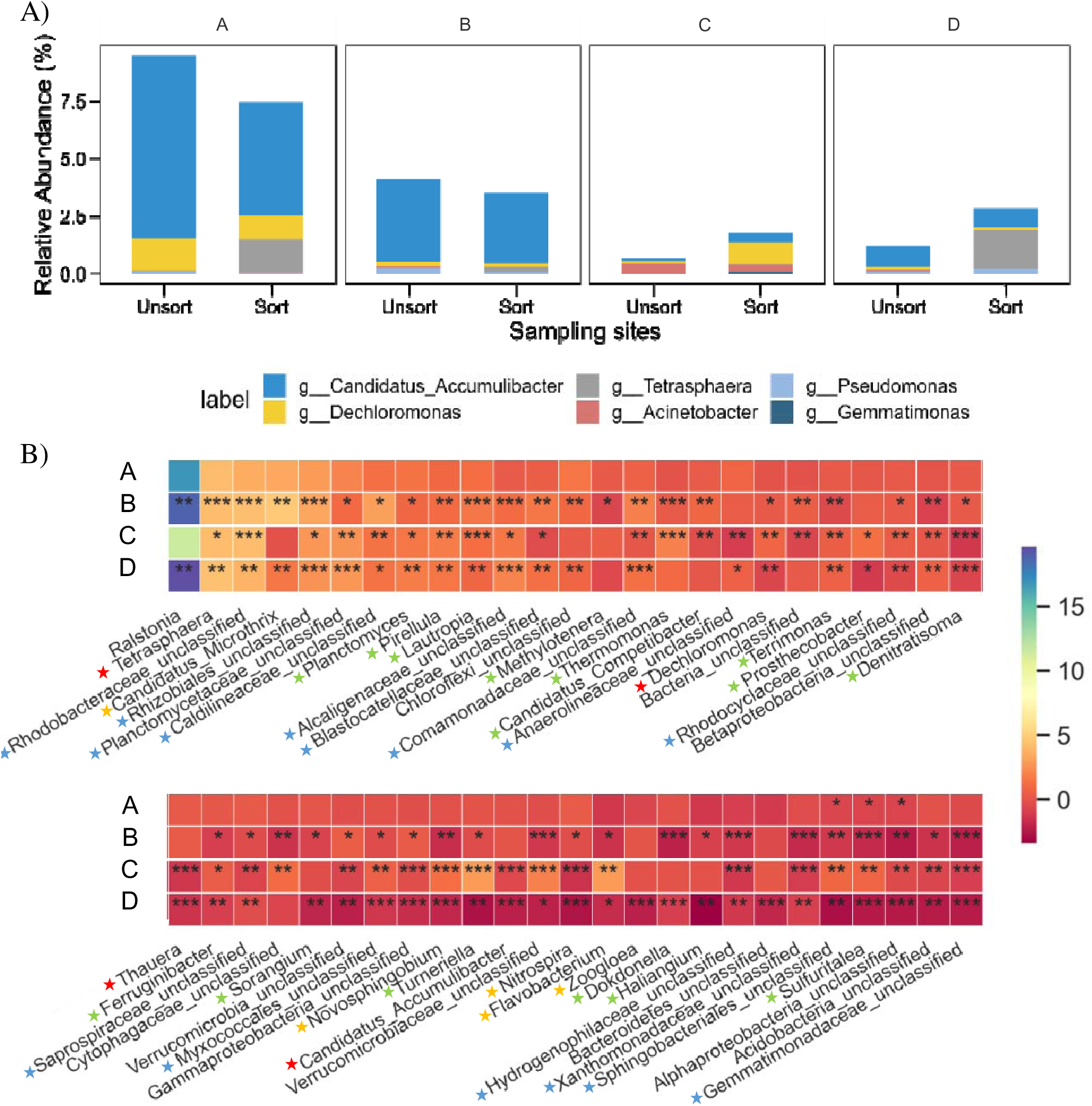
A) Relative abundances of known candidate PHA-PAOs before (unsort) and after (sort) TriFlow-seq in three full-scale EBPR plants and one pilot-scale S2EBPR plant. Groups with relative abudances below 0.1% are not shown; B) Differential abundance heatmap of bacterial genera before and after TriFlow-seq sorting. The heat map represents t-statistics derived from two-tailed t-tests on log_10_-transformed fold changes in relative abundance of bacterial genera after TriFlow sorting in comprison to prior-sorting. Blue hues denote positive t-statistics (enrichment via sorting), while red hues indicate negative t-statistics (depletion after sorting). Statistical significance levels: \**p* < 0.05, \*\**p* < 0.01, \*\*\**p* < 0.001 The red pentagram indicates the genus has been experimentally confirmed with PHA/polyP and PHA-PAO activities; the yellow pentagram indicates the presence of polyP and PHA; the green pentagram indicates that the reference genome contains necessary genes for polyP and carbon storage synthesis; the blue pentagram indicates that some genera within the Family have polyP accumulation ability. Detailed information can be found in Table S3. Analysis is limited to the top 50 genera by abundance.

**Figure 4.**
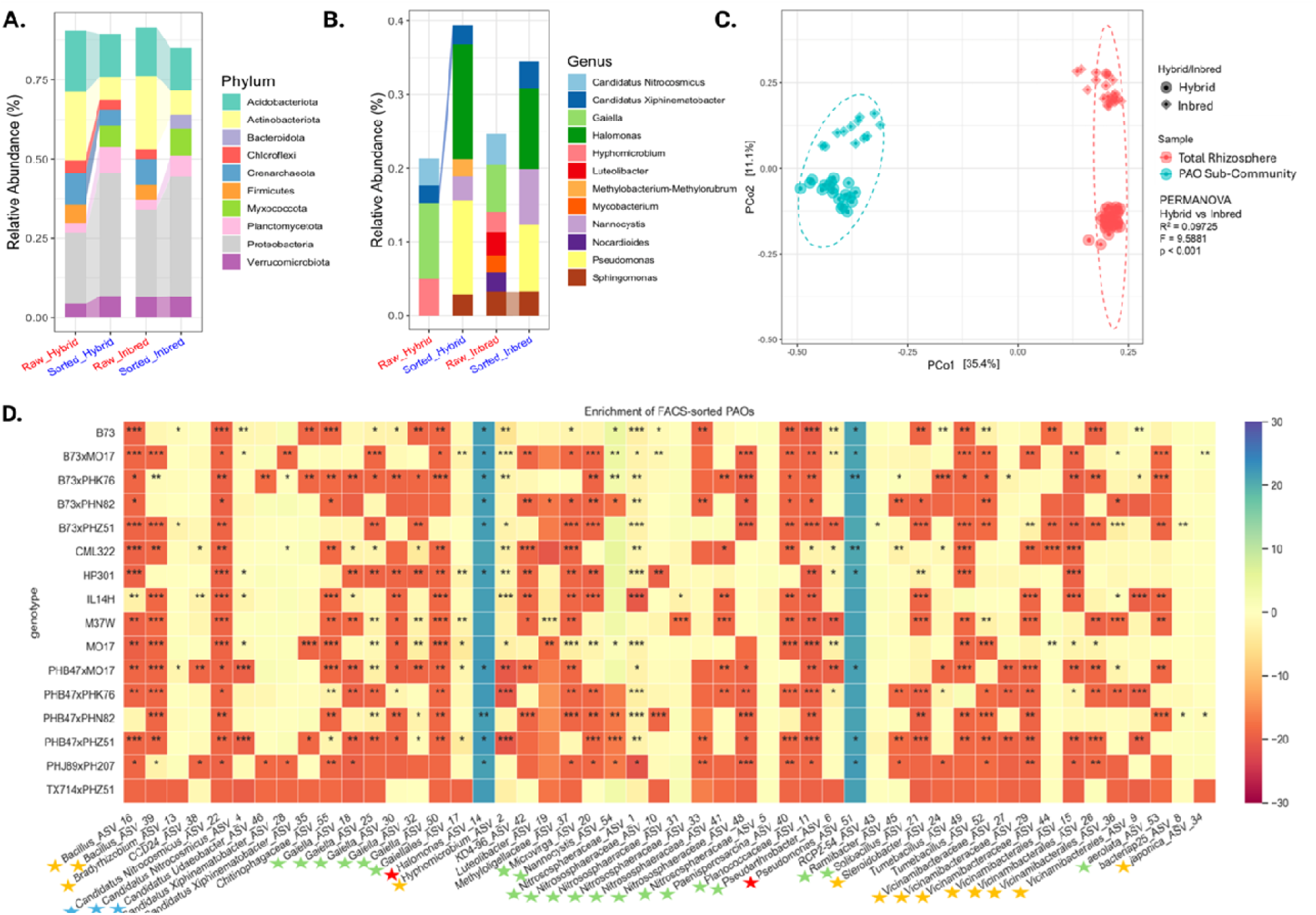
of candidate rhizosphere PHA-PAOs and characterization of the rhizosphere PHA-PAO community vi TriFlow-Seq. A) Phylum- and B) Genus-level community composition and relative abundance of unsorted raw rhizosphere community and sorted PHA-PAO community, separated by field experiment, abundance filtered > 0.03; B) Principal coordinate analysis (PCoA) based on Bray-Curtis dissimilarity of raw rhizosphere community an FACS-sorted PHA-PAO community across maize inbred and hybrid lines; D) Differential abundance analysis of candidate rhizosphere PHA-PAO taxa in inbred (e.g., B73) and hybrid (e.g., B73×Mo17) rhizospheres following TriFlow-seq. The blue scale denotes positive enrichment in the PHA-PAO community, and the red scale denotes negative enrichment. Significance is based on a two-tailed t-test with an unscaled log_10_-fold change: \**p* < 0.05, \*\**p* < 0.01, \*\*\**p* < 0.001. The red pentagram indicates the genus has been experimentally confirmed with PHA/polyP and PHA-PAO activities; the yellow pentagram indicates the presence of polyP and PHA; the green pentagram indicates that the reference genome contains necessary genes for polyP and carbon storage synthesis; the blue pentagram indicates that some genera within the Family have polyP accumulation ability. Detailed information can be found in Table S4. Analysis was limited to the top 50 amplicon sequence variants (ASVs) by abundance.

### 2.10 Single cell Raman spectroscopy (SCRS)

As a second validation, we compared FACS results with SCRS analysis, an independent method previously developed by our group to quantify polyP- and PHA-containing cells.^36,37^ Details of SCRS analysis for PHA-PAO identification, polyP and PHA detection can be referred to our previous publications.^58^ Key details are provided in Text S5.

### 2.11 Statistical analysis

Spearman and Pearson correlation coefficients and corresponding *p*-values were computed using the R *stats* (4.1.3) package. Principal coordinates analysis (PCoA) was performed using the R *vegan* (2.6-4) package to assess beta diversity based on Bray-Curtis dissimilarities of normalized relative abundance data. The ordination plot was visualized using the R *ggplot2* (3.4.4) package. For the differential abundance analysis, log -transformed fold changes were compared between genera using two-tailed independent t-tests implemented in Python (version 3.12) with SciPy (version 1.10.1). The resulting t-statistics were visualized as a heatmap using the Seaborn package (version 0.13.2).

## 3 Results and discussion

### 3.1 Evaluation of TriFlow for sorting PHA-containing PAOs

The effectiveness of TriFlow for identifying and sorting PHA-containing PAOs was tested with sludge samples from EBPR and S2EBPR systems, which are known to be enriched in PHA- PAOs and served as positive environmental controls. The density of the triple-stained cells before and after the TriFlow enrichment was visualized using confocal microscopy (Figure 1A). A significant and evident enrichment of candidate PHA-PAO cells containing both polyP and PHA was observed in the sorted samples (magenta) compared to the original sludge, a pattern validated in all four tested EBPR systems. In addition, the relative abundances of PHA-positive and polyP-positive cells to total cells (SG-stained cells) were semi-quantified using digital image analysis (Figure 1B). NR-stained and TC-stained cells were clearly enriched in all FACS sorted samples compared to the original sludge samples. These results demonstrate that our gating strategy in the FACS analysis effectively enriched dual polyP- and PHA-accumulating PAOs as expected, confirming the appropriateness of the gating strategy employed.

### 3.2 Validation of PHA-PAOs quantification using TriFlow

To Further validate the TriFlow method, we compared the relative abundance of PHA-PAOs identified through FACS with those determined by SCRS across various microbial samples, including both EBPR systems biomass and the maize rhizosphere soil samples (Figure 2). SCRS, which targets the molecular composition of microbial cells, has demonstrated exceptional sensitivity and accuracy in detecting and quantifying intracellular polymers such as polyP, PHA, and glycogen.^38^ SCRS-based detection of PHA-PAOs containing both polyP and PHA has been demonstrated in previous studies^27,36–39^ Importantly, the percentage of PHA-PAOs determined by FACS and SCRS fell within the same range. A significant positive monotonic relationship was observed between the two techniques (Spearman’s ρ = 0.62, *p* = 0.009), demonstrating statistically robust concordance in detecting functional PHA-PAOs by these two independent methods.

Although the percentages of PHA-PAOs determined by TriFlow were consistent and correlated with those identified by SCRS, discrepancies occurred due to inherent differences in detection principles and sample processing protocols. SCRS is a non-invasive, fixation-free method that detects intracellular PHA and polyP based on their unique Raman spectral signatures and has proven effective in detecting multiple intracellular polymers with relatively less influence from varying polyP composition (i.e., counter metal ions).^59,60^ In contrast, FACS requires cell fixation and permeation steps to enhance dye penetration, which permanently alters cell wall and membrane permeability.^47^ This modification typically results in a 10-20% loss of intracellular polymers, similar to observations with fluorescence *in situ* hybridization (FISH) fixatives.^39^ In addition, the accuracy of FACS largely relies on the specificity and sensitivity of fluorescent dyes. Nile red, although widely employed for PHA detection in flow cytometry, can also bind to other lipophilic inclusions such as trehalose and triglycerides.^61^ Similarly, tetracycline detects polyP by preferentially complexing with countercharged divalent cations (e.g., Ca², Mg²) in polyP granules.^35,62,63^ These methodological differences suggest that FACS may slightly overestimate PHA-containing PHA-PAOs relative to the SCRS method, consequently reducing the slope of the linear curve. Furthermore, FACS employs size-based filtration (40 μm cell strainer) to prevent flow cytometer nozzle clogging, while SCRS utilizes density-gradient centrifugation for cell extraction.^6460^ These procedural variations likely influence the organisms being analyzed, contributing to the observed differences in PHA-PAO abundance percentages. Despite these discrepancies, the consistent trend in PHA-PAO detection by both methods demonstrates the viability of the TriFlow for identifying and quantifying PHA- PAOs in complex microbial communities.

### 3.3 Application of TriFlow-Seq for the identification and quantification of PHA-PAOs in EBPR systems

Combining TriFlow with downstream 16S rRNA gene amplicon sequencing (termed TriFlow- Seq) enabled the enrichment, quantification, and identification of both known and candidate PHA-PAOs in EBPR systems. The relative abundances of canonical PHA-PAOs in the original sludge prior to cell sorting and after TriFlow enrichment in pilot- and full-scale EBPR plants are depicted in Figure 3.

#### 3.3.1 Enrichment of known PHA-PAOs in EBPR systems using TriFlow-Seq

Across all sampled wastewater treatment plants, TriFlow-Seq consistently detected well-established PHA-PAO genera, including *Pseudomonas*, *Dechloromonas*, and *Ca. Accumulibacter* (Figure 3A). These genera are recognized for their dual accumulation of polyP and PHA in EBPR systems,^65–68^ validating TriFlow-Seq’s ability to target functionally active PHA-PAOs.

To further characterize the sorted communities, we calculated enrichment ratios, defined as the log_10_-transformed ratio of post-sorting to pre-sorting relative abundances for each genus. Higher ratios indicate a greater enrichment of bacteria capable of accumulating both polyP and intracellular C polymers, thus reflecting the proportion of putative PHA-PAOs within that taxonomic group (Figure 3B). It should be noted that even within known PHA-PAOs, some sub- genus populations were not actively performing polyP and PHA accumulation, resulting in a negative enrichment factor, as observed for *Ca*. Accumulibacter. Despite being consistently identified as the dominant PHA-PAO in raw sludge samples from the three full-scale plants (except for the S2EBPR pilot plant, with 1.2∼4.0%) (Figure 3A), *Ca*. Accumulibacter showed varied enrichment ratios among the EBPR plants (Figure 3B). Its relative abundance even decreased after sorting in sample C. Interestingly, *Tetrasphaera* emerged as the most significantly enriched genus after sorting (Figure 3A and 3B). Although initially thought to lack PHA synthesis capability, recent studies have revealed that anaerobic PHA synthesis depends on clade composition of *Tetrasphaera*.^11–14^ For instance, the *T. japonica* genome possesses PHA synthase (*phaC*) and has been experimentally shown to synthesize PHA under anaerobic conditions.^11^ This evidence suggests that *T. japonica*, or potentially uncultured *Tetrasphaera* with Accumulibacter-like PAO metabolism, resides in full-scale EBPR systems.^69^ Overall, the relative abundances of known PHA-PAOs accounted for only 2 to 12% of total PHA+polyP-positive cells (Figure 3A), highlighting the greater phylogenetic diversity of PHA-PAOs than our current understanding.

#### 3.3.2 Discovery of candidate PHA-PAOs in EBPR systems

Based on TriFlow results, we proposed putative PHA-PAOs in EBPRs based on (1) consistent detection across sorted communities via TripleFlow-seq; (2) genomic potential for polyP and PHA accumulation within the detected genus. A diverse putative PHA-PAOs, including unclassified Rhodobacteraceae, unclassified Rhizobiales, unclassified Caldilineaceae, *Lautropia*, and others (Figure 3B). Among them, a substantial portion lacked genus-level classification, suggesting the presence of uncharacterized genera that may possess PHA-PAO-like functions. Although these families or orders have not traditionally been associated with PHA-PAO activity, related members within the same taxonomic groups have been reported to accumulate intracellular polyP and PHA based on genomics analysis. For instance, some bacteria isolated from the order Rhizobiales were found to contain PHA and polyP.^70^ An uncultured genus close to the Rhodobacteraceae family is speculated to be sulfur PHA-PAOs.^71^ These findings provide preliminary evidence for novel candidate PHA-PAOs, though further studies are needed to determine whether they contribute to phosphorus removal through classical or alternative EBPR pathways.

These results revealed important insights into PHA-PAO diversity and functionality in EBPR systems. Not all genetically identified *Ca*. Accumulibacter displayed metabolically active PAO phenotypes, as evidenced by their varied enrichment ratios, highlighting the distinction between genetic potential and phenotypic expression. *Tetrasphaera* showed unexpected and significant enrichment with variable intracellular PHA content, revealing phenotypic diversity beyond the current understanding of this genus. Additionally, the TriFlow-Seq method uncovered novel candidate PHA-PAOs from previously unrecognized taxonomic groups, expanding our understanding of PHA-PAO diversity in EBPR systems. Although further investigation and confirmation of the roles of these candidate PHA-PAOs in the tested plants is beyond the focus of this study, this application has demonstrated that TriFlow-Seq is a valuable tool for rapidly enriching and identifying targeted bacterial populations that exhibit the phenotypic traits characteristic of functionally relevant PHA-PAOs. This approach was further adapted and optimized to be applicable to more heterogeneous sample matrices, such as bulk and rhizosphere soil, enabling the identification and characterization of PHA-PAOs in these complex environmental matrices.

### 3.4 Discovery of the identities of previously unknown rhizosphere PHA-PAOs communities

#### 3.4.1 Effectiveness in application of TriFlow-Seq in rhizosphere samples

Application of targeted single-cell FACS for PHA-PAO identification in soil samples has not been previously reported. Our newly developed TriFlow-Seq pretreatment method and sequential targeted gating successfully identified and isolated PHA-PAO populations from the rhizosphere microbiome. Genomic DNA recovery ranged from 1 ng/µL to 16.3 ng/µL, with high-quality (Phred >30) 16S rRNA gene amplicon reads obtained in all but one sequenced sorted sample (*n* = 192), demonstrating the method’s effectiveness across diverse soil types.

Successful application and recovery of the 16S rRNA gene amplicon enabled taxonomic identification of the previously unknown PHA-PAOs community, revealed to include the unanimous presence of canonical PHA-PAOs associated EBPR systems and performance. Proteobacteria, Actinobacteria, and Acidobacteria were the most abundant phyla in both unsorted and sorted communities (63.2-66.2% and 59.1-59.5% respectively) (Figure 4A), aligning with the typically observed community compositions in both maize rhizosphere and BEPR communities.^72^ Notably, the phylum Myxococcota, known for complex social behaviors and recently identified to harbor *ppk1* associated with polyP accumulation in activated sludge,^73,74^ was consistently detected only within the sorted communities (6.58-8.54%) (Figure 4A). At the genus level, while typical PHA-PAOs such as *Tetrasphaera*, *Ca.* Accumulibacter, and *Dechloromonas* were detected within the sorted community, their combined abundance was below 1%, suggesting minimal contribution to the rhizosphere processes. Instead, *Pseudomonas* (9.03-12.7%) and *Halomonas* (11-15.6%), both putative PHA-PAOs identified in EBPR systems, were the predominant genera in the sorted communities (Figure 4B). Further, of the conventionally known PHA-PAOs, including *Ca. Accumulibacter*, *Tetrasphaera*, *Dechloromonas*, *Acinetobacter*, *Gemmatimonas*, and *Pseudomonas*. there was no significant loss of unrarefied observed species richness within the *Tetrasphaera* and *Acinetobacter* genera between the sorted and unsorted communities indicating the scope and accuracy of our TriFlow- Seq method in surveying and conserving the active PHA-PAO diversity in rhizosphere samples; capable of identifying functionally verified PHA-PAOs despite naturally low abundances (Figure S7).

#### 3.4.2 Discovery of novel PHA-PAOs in the maize rhizosphere

In addition to the discovery of canonical PHA-PAO activity in the rhizosphere, for the first time we were able to capture a markedly different co-polymer accumulating community than those reported in EBPR systems and propose novel candidate PHA-PAOs functionally active in rhizosphere systems. The enrichment profiles of key sorted taxa across different maize genotypes were examined to identify candidate functional PHA-PAOs in the maize rhizosphere. The selection criteria used were comparable to those established for activated sludge systems in Section 3.3.1 but adapted to consider the inherent variability of soil systems which leads to differences in phenotypic profiles and diverse metabolic activities among potential PHA-PAOs.

The enrichment profiles at the sub-genus level revealed the selective dominance of specific amplicon sequence variants (ASVs) within the predominate sorted genera, leading to the identification of three candidate rhizosphere PHA-PAOs. Among these, two predominant putative PHA-PAO ASVs exhibited positive enrichment across all maize lines: *Halomonas* ASV_14 (21.1-22.1-fold) and *Pseudomonas* ASV_51 (20.7-22-fold) (Figure 4D). Additionally, *Nannocystis* ASV_54, within the Myxococcota phylum, most closely related to *Nannocystis exedens* (93.68% 16S rRNA sequence similarity, BLASTN), exhibited positive enrichment (1.14-4.46-fold) across 68.75% of samples, specifically in sampled inbreds (including B73) and most maternal B73 hybrids (*p*-value <0.05) (Figure 4D). Analysis of the available annotated genome of *N. exedens* (GenBank GCA_002343915.1) revealed the presence of *ppk1*, *ppx*, and sterol and steroid biosynthetic pathways.^75^ While this study does not aim to exhaustively compare PHA-PAO populations across diverse maize genotypes, our proof-of-concept analysis revealed distinct identities of novel PHA-PAO with distinct key C and P functions and their unique associations with different maize genotypes, warranting further investigation.

#### 3.4.3 Exploration of rhizosphere PHA-PAO community dynamics

In addition to revealing the PHA-PAO community’s taxonomic identity, we first investigated community dynamics across maize host genetic diversity, an emerging driver in rhizosphere structure.^72,76^ Principal coordinate analysis (PCoA) based on Bray-Curtis dissimilarity revealed distinct clustering and significant separation between the unsorted and sorted community along PCo1 (35.4% variance; PERMANOVA, *p*-value <0.001), indicating isolation of a distinct sub- community (Figure 4C). Within both clusters, two significant sub-clusters emerged based on maize genetic background (inbred versus hybrid) along PCo2 (11.1% variation, *p*-value <0.001, PERMANOVA) (Figure 4C), suggesting that the sorted population retains its community fingerprint from its original rhizosphere population. Alpha diversity was significantly reduced in sorted samples across all metrics, including species richness, Shannon index, and Simpson’s index (*p* < 0.001; Figure S10), confirming that TriFlow-Seq enriched a specialized sub- community dominated by a few taxa.

These results, for the first time, revealed the characteristics, diversity, composition, and distribution pattern of the rhizosphere PHA-PAO community. Emerging evidence highlighted the important roles of PHA-PAOs in rhizosphere communities and processes, including halotolerance, phytohormone production, siderophore production, beneficial antagonism/antibiotic production, and nutrient cycling.^77–79^ Our TriFlow-Seq method enabled the discovery of new PHA-PAOs in agricultural soils, expanding our understanding of their identity and diversity within agricultural systems.

## 4 TriFlow-Seq: Achievements, Applications, and Future Prospects

Linking phenotype-based detection and genotype-based characterization provides valuable insights into the discovery of organisms of interest. In this study, we developed and validated the TriFlow-Seq workflow and demonstrated two key applications: identifying functional PHA- PAOs in wastewater treatment facilities and uncovering previously unrecognized PHA-PAOs involved in P and C cycling within the maize rhizosphere. By modifying FACS preprocessing protocols, we enabled the identification of candidate PHA-PAOs in modified environments, such as soil matrix. Our findings revealed unexpectedly high PHA-PAO prevalence and distinct phylogenetic patterns across different maize genotypes, establishing a pioneering approach to exploring PHA-PAO identities and functions in rhizosphere systems.

Future applications of Reflow-Seq can be enhanced through several approaches: (1) incorporating additional phenotyping targets as new staining materials become available, potentially revealing greater PHA-PAO diversity; (2) integrating mini-metagenomics or other omics techniques to obtain deeper functional and metabolic insights into candidate PHA-PAOs; and (3) combining with FISH to increase specificity for targeting particular microbial groups. During our investigation, tetracycline and DAPI were the only fluorescent dyes proven effective for FACS-based PHA-PAO selection.^34,35^ However, recent advancements have introduced more specific dyes, such as JC-D7, which offer improved specificity for polyP detection in environmental samples.^80^ Incorporating such novel dyes into the TriFlow-Seq workflow could enhance the accuracy and sensitivity of PHA-PAO identification in future research. Despite current limitations, this technique demonstrates significant potential for identifying novel PHA- PAOs within complex microbial consortia, advancing our understanding of microbial polyP metabolism and phosphorus cycling across diverse ecosystems.

## Acknowledgements

This study was partially supported by the Cornell Institute for Digital Agriculture (CIDA), and the Center for Research on Programmable Plant System (CROPPS) funded by the National Science Foundation (Grant No. DBI-2019674). We thank Dr. Lydia Tesfa, Jaclyn Mahoney, and Michael Sledziona from the Flow Cytometry Facility of the Biotechnology Resource Center at Cornell University (RRID:SCR_021740) for their technical assistance with FACS experiments. The authors would like to thank the G2F consortium for providing the High-Intensity Phenotyping Site (HIPS) experiment, which consisted of hybrids that were used in this study. This consortium involves more than 30 researchers representing more than 20 research institutions. Details about the initiative and publicly available resources can be found at www.Genomes2Fields.org. We thank Dr. Maria Cinta Romay and Nicholas Lepak for their generous time and effort in coordinating and planting the HIPS experiment.

---

**For Table of Contents Only**

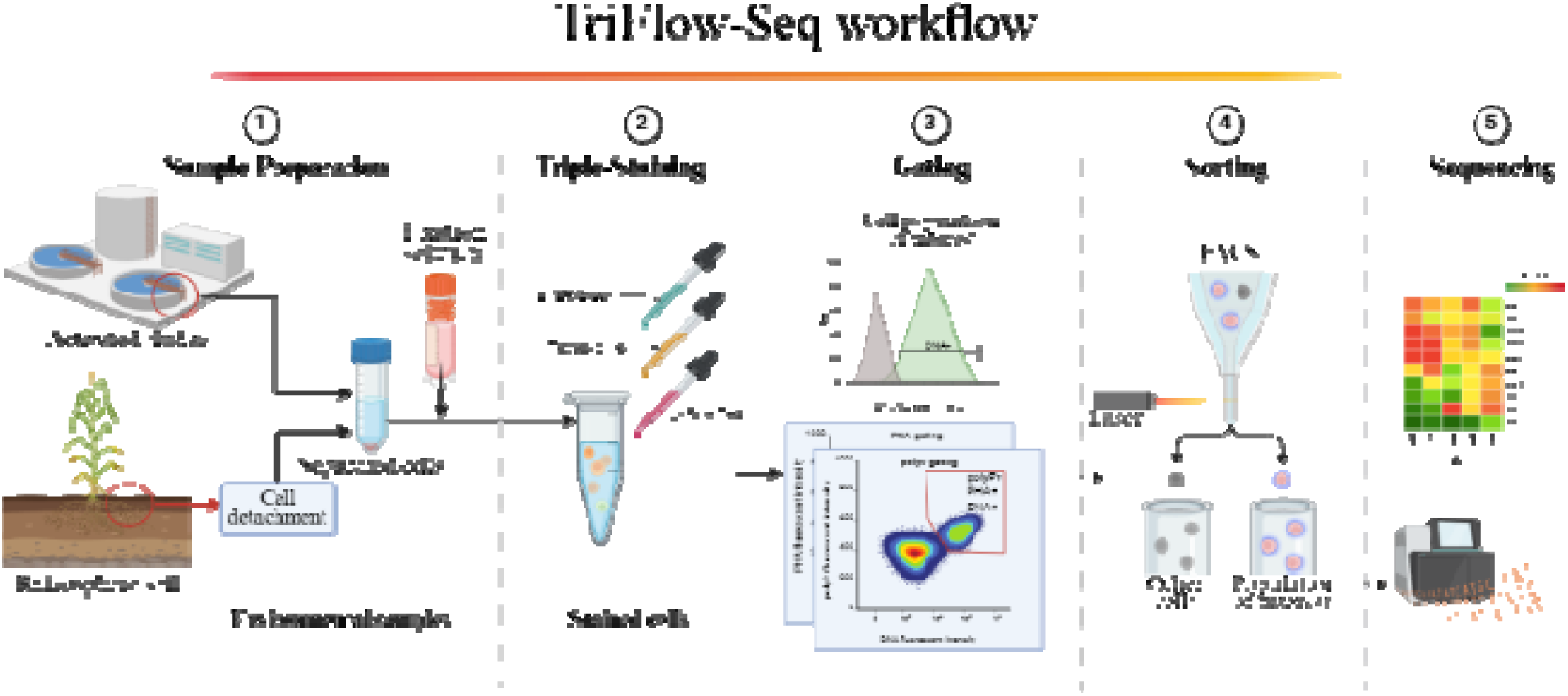

## References

(1) Mason-Jones, K.; Robinson, S. L.; Veen, G. F. (Ciska); Manzoni, S.; van der Putten, W. H. Microbial Storage and Its Implications for Soil Ecology. ISME J 2022, 16 (3), 617–629. 10.1038/s41396-021-01110-w.

(2) Rao, N. N.; Gómez-García, M. R.; Kornberg, A. Inorganic Polyphosphate: Essential for Growth and Survival. Annual Review of Biochemistry 2009, 78 (Volume 78, 2009), 605–647. 10.1146/annurev.biochem.77.083007.093039.

(3) Kornberg, A.; Rao, N. N.; Ault-Riché, D. Inorganic Polyphosphate: A Molecule of Many Functions. Annual Review of Biochemistry 1999, 68 (1), 89–125. 10.1146/annurev.biochem.68.1.89.

(4) Oehmen, A.; Lemos, P. C.; Carvalho, G.; Yuan, Z.; Keller, J.; Blackall, L. L.; Reis, M. A. M. Advances in Enhanced Biological Phosphorus Removal: From Micro to Macro Scale. Water Research 2007, 41 (11), 2271–2300. 10.1016/j.watres.2007.02.030.

(5) Seviour, R. J.; Mino, T.; Onuki, M. The Microbiology of Biological Phosphorus Removal in Activated Sludge Systems. FEMS Microbiology Reviews 2003, 27 (1), 99–127. 10.1016/S0168-6445(03)00021-4.

(6) Martín, H. G.; Ivanova, N.; Kunin, V.; Warnecke, F.; Barry, K. W.; McHardy, A. C.; Yeates, C.; He, S.; Salamov, A. A.; Szeto, E.; Dalin, E.; Putnam, N. H.; Shapiro, H. J.; Pangilinan, J. L.; Rigoutsos, I.; Kyrpides, N. C.; Blackall, L. L.; McMahon, K. D.; Hugenholtz, P. Metagenomic Analysis of Two Enhanced Biological Phosphorus Removal (EBPR) Sludge Communities. Nat Biotechnol 2006, 24 (10), 1263–1269. 10/dct6rr.

(7) Petriglieri, F.; Singleton, C.; Peces, M.; Petersen, J. F.; Nierychlo, M.; Nielsen, P. H. “Candidatus Dechloromonas Phosphoritropha” and “Ca. D. Phosphorivorans”, Novel Polyphosphate Accumulating Organisms Abundant in Wastewater Treatment Systems. ISME J 2021, 15 (12), 3605–3614. 10.1038/s41396-021-01029-2.

(8) Li, G.; Tooker, N. B.; Wang, D.; Srinivasan, V.; Barnard, J. L.; Russell, A.; Stinson, B.; McQuarrie, J.; Schauer, P.; Menniti, A.; Varga, E.; Hauduc, H.; Takács, I.; Bott, C.; Dobrowski, P.; Onnis-Hayden, A.; Gu, A. Z. Modeling Versatile and Dynamic Anaerobic Metabolism for PAOs/GAOs Competition Using Agent-Based Model and Verification via Single Cell Raman Micro-Spectroscopy. Water Research 2023, 245, 120540. 10/gs5c3w.

(9) Zhao, W.; Bi, X.; Peng, Y.; Bai, M. Research Advances of the Phosphorus-Accumulating Organisms of Candidatus Accumulibacter, Dechloromonas and Tetrasphaera: Metabolic Mechanisms, Applications and Influencing Factors. Chemosphere 2022, 307, 135675. 10/gq8m45.

(10) Tobin, K. M.; McGrath, J. W.; Mullan, A.; Quinn, J. P.; O’Connor, K. E. Polyphosphate Accumulation by Pseudomonas Putida CA-3 and Other Medium-Chain-Length Polyhydroxyalkanoate-Accumulating Bacteria under Aerobic Growth Conditions. Applied and Environmental Microbiology 2007, 73 (4), 1383–1387. 10.1128/AEM.02007-06.

(11) Kristiansen, R.; Nguyen, H. T. T.; Saunders, A. M.; Nielsen, J. L.; Wimmer, R.; Le, V. Q.; McIlroy, S. J.; Petrovski, S.; Seviour, R. J.; Calteau, A.; Nielsen, K. L.; Nielsen, P. H. A Metabolic Model for Members of the Genus Tetrasphaera Involved in Enhanced Biological Phosphorus Removal. ISME J 2013, 7 (3), 543–554. 10.1038/ismej.2012.136.

(12) Close, K.; Marques, R.; Carvalho, V. C. F.; Freitas, E. B.; Reis, M. A. M.; Carvalho, G.; Oehmen, A. The Storage Compounds Associated with Tetrasphaera PAO Metabolism and the Relationship between Diversity and P Removal. Water Research 2021, 204, 117621. 10/gsmsnj.

(13) Marques, R.; Santos, J.; Nguyen, H.; Carvalho, G.; Noronha, J. P.; Nielsen, P. H.; Reis, M. A. M.; Oehmen, A. Metabolism and Ecological Niche of Tetrasphaera and Ca. Accumulibacter in Enhanced Biological Phosphorus Removal. Water Research 2017, 122, 159–171. 10.1016/j.watres.2017.04.072.

(14) McIlroy, S. J.; Onetto, C. A.; McIlroy, B.; Herbst, F. A.; Dueholm, M. S.; Kirkegaard, R. H.; Fernando, E.; Karst, S. M.; Nierychlo, M.; Kristensen, J. M.; Eales, K. L.; Grbin, P. R.; Wimmer, R.; Nielsen, P. H. Genomic and in Situ Analyses Reveal the Micropruina Spp. as Abundant Fermentative Glycogen Accumulating Organisms in Enhanced Biological Phosphorus Removal Systems. Frontiers in Microbiology 2018, 9 (MAY), 1–12. 10.3389/fmicb.2018.01004.

(15) Akbari, A.; Wang, Z.; He, P.; Wang, D.; Lee, J.; Han, I.; Li, G.; Gu, A. Z. Unrevealed Roles of Polyphosphate accumulating Microorganisms. Microb Biotechnol 2021, 14 (1), 82–87. 10.1111/1751-7915.13730.

(16) Cordell, D.; Rosemarin, A.; Schröder, J. J.; Smit, A. L. Towards Global Phosphorus Security: A Systems Framework for Phosphorus Recovery and Reuse Options. Chemosphere 2011, 84 (6), 747–758. 10/d4mvgw.

(17) Zabel, F.; Putzenlechner, B.; Mauser, W. Global Agricultural Land Resources–a High Resolution Suitability Evaluation and Its Perspectives until 2100 under Climate Change Conditions. PloS one 2014, 9 (9), e107522.

(18) Saia, S. M.; Carrick, H. J.; Buda, A. R.; Regan, J. M.; Walter, M. T. Critical Review of Polyphosphate and Polyphosphate Accumulating Organisms for Agricultural Water Quality Management. Environ. Sci. Technol. 2021, 55 (5), 2722–2742. 10.1021/acs.est.0c03566.

(19) Mason-Jones, K.; Banfield, C. C.; Dippold, M. A. Compound-Specific 13C Stable Isotope Probing Confirms Synthesis of Polyhydroxybutyrate by Soil Bacteria. Rapid Communications in Mass Spectrometry 2019, 33 (8), 795–802. 10/gr8ctq.

(20) Mason-Jones, K.; Breidenbach, A.; Dyckmans, J.; Banfield, C. C.; Dippold, M. A. Intracellular Carbon Storage by Microorganisms Is an Overlooked Pathway of Biomass Growth. Nat Commun 2023, 14 (1), 2240. 10/gr745w.

(21) Butler, O. M.; Manzoni, S.; Warren, C. R. Community Composition and Physiological Plasticity Control Microbial Carbon Storage across Natural and Experimental Soil Fertility Gradients. ISME J 2023, 17 (12), 2259–2269. 10.1038/s41396-023-01527-5.

(22) Ratcliff, W. C.; Kadam, S. V.; Denison, R. F. Poly-3-Hydroxybutyrate (PHB) Supports Survival and Reproduction in Starving Rhizobia: PHB Increases Rhizobium Fitness. FEMS Microbiology Ecology 2008, 65 (3), 391–399. 10.1111/j.1574-6941.2008.00544.x.

(23) Smith, S. E.; Smith, F. A. Roles of Arbuscular Mycorrhizas in Plant Nutrition and Growth: New Paradigms from Cellular to Ecosystem Scales. Annual Review of Plant Biology 2011, 62 (Volume 62, 2011), 227–250. 10.1146/annurev-arplant-042110-103846.

(24) Rao, N.; Kornberg, A. Inorganic Polyphosphate Regulates Responses of Escherichia Coli to Nutritional Stringencies, Environmental Stresses and Survival in the Stationary Phase. Inorganic polyphosphates: Biochemistry, biology, biotechnology 1999, 183–195.

(25) Gray, M. J.; Jakob, U. Oxidative Stress Protection by Polyphosphate — New Roles for an Old Player. Current Opinion in Microbiology 2015, 24, 1–6. 10.1016/j.mib.2014.12.004.

(26) Alam, M. M.; Srinivasan, V.; Mueller, A. V.; Gu, A. Z. Status and Advances in Technologies for Phosphorus Species Detection and Characterization in Natural Environment- A Comprehensive Review. Talanta 2021, 233, 122458. 10.1016/j.talanta.2021.122458.

(27) Li, Y.; Cope, H. A.; Rahman, S. M.; Li, G.; Nielsen, P. H.; Elfick, A.; Gu, A. Z. Toward Better Understanding of EBPR Systems via Linking Raman-Based Phenotypic Profiling with Phylogenetic Diversity. Environmental Science and Technology 2018, 52 (15), 8596–8606. 10.1021/acs.est.8b01388.

(28) Majed, N.; Li, Y.; Gu, A. Z. Advances in Techniques for Phosphorus Analysis in Biological Sources. Current Opinion in Biotechnology 2012, 23 (6), 852–859. 10.1016/j.copbio.2012.06.002.

(29) He, Z.; Deng, Y.; Van Nostrand, J. D.; Tu, Q.; Xu, M.; Hemme, C. L.; Li, X.; Wu, L.; Gentry, T. J.; Yin, Y.; Liebich, J.; Hazen, T. C.; Zhou, J. GeoChip 3.0 as a High-Throughput Tool for Analyzing Microbial Community Composition, Structure and Functional Activity. The ISME Journal 2010, 4 (9), 1167–1179. 10.1038/ismej.2010.46.

(30) Tu, Q.; Yu, H.; He, Z.; Deng, Y.; Wu, L.; Van Nostrand, J. D.; Zhou, A.; Voordeckers, J.; Lee, Y.; Qin, Y. GeoChip 4: A Functional Gene array based High throughput Environmental Technology for Microbial Community Analysis. Molecular ecology resources 2014, 14 (5), 914–928.

(31) Kornberg, A. Inorganic Polyphosphate: Toward Making a Forgotten Polymer Unforgettable. Journal of Bacteriology 1995, 177 (3), 491–496. 10.1128/jb.177.3.491-496.1995.

(32) Kulaev, I. S.; Vagabov, V.; Kulakovskaya, T. The Biochemistry of Inorganic Polyphosphates, 2 nd.; John Wiley & Sons, 2005.

(33) Whitehead, M. P.; Eagles, L.; Hooley, P.; Brown, M. R. W. Most Bacteria Synthesize Polyphosphate by Unknown Mechanisms. Microbiology 2014, 160 (5), 829–831. 10.1099/mic.0.075366-0.

(34) Terashima, M.; Kamagata, Y.; Kato, S. Rapid Enrichment and Isolation of Polyphosphate-Accumulating Organisms Through 4’6-Diamidino-2-Phenylindole (DAPI) Staining With Fluorescence-Activated Cell Sorting (FACS). Front. Microbiol. 2020, 11. 10.3389/fmicb.2020.00793.

(35) Günther, S.; Trutnau, M.; Kleinsteuber, S.; Hause, G.; Bley, T.; Röske, I.; Harms, H.; Müller, S.; Gunther, S.; Kleinsteuber, S.; Trutnau, M.; Bley, T.; Muller, S.; Hause, G.; Roske, I.; Harms, H.; Gunther, S.; Bley, T.; Muller, S.; Hause, G.; Roske, I.; Harms, H. Dynamics of Polyphosphate-Accumulating Bacteria in Wastewater Treatment Plant Microbial Communities Detected via DAPI (4′,6′-Diamidino-2- Phenylindole) and Tetracycline Labeling. Applied and Environmental Microbiology 2009, 75 (7), 2111–2121. 10.1128/AEM.01540-08.

(36) Wang, D.; He, P.; Wang, Z.; Li, G.; Majed, N.; Gu, A. Z. Advances in Single Cell Raman Spectroscopy Technologies for Biological and Environmental Applications. Current Opinion in Biotechnology 2020, 64, 218–229. 10.1016/j.copbio.2020.06.011.

(37) Majed, N.; Chernenko, T.; Diem, M.; Gu, A. Z. Identification of Functionally Relevant Populations in Enhanced Biological Phosphorus Removal Processes Based on Intracellular Polymers Profiles and Insights into the Metabolic Diversity and Heterogeneity. Environmental Science and Technology 2012, 46 (9), 5010–5017. 10.1021/es300044h.

(38) Majed, N.; Matthäus, C.; Diem, M.; Gu, A. Z. Evaluation of Intracellular Polyphosphate Dynamics in Enhanced Biological Phosphorus Removal Process Using Raman Microscopy. Environmental Science and Technology 2009, 43 (14), 5436–5442. 10.1021/es900251n.

(39) Fernando, E. Y.; McIlroy, S. J.; Nierychlo, M.; Herbst, F.-A.; Petriglieri, F.; Schmid, M. C.; Wagner, M.; Nielsen, J. L.; Nielsen, P. H. Resolving the Individual Contribution of Key Microbial Populations to Enhanced Biological Phosphorus Removal with Raman–FISH. ISME J 2019, 13 (8), 1933–1946. 10/gn7s5g.

(40) McMullen, M. D.; Kresovich, S.; Villeda, H. S.; Bradbury, P.; Li, H.; Sun, Q.; Flint-Garcia, S.; Thornsberry, J.; Acharya, C.; Bottoms, C.; Brown, P.; Browne, C.; Eller, M.; Guill, K.; Harjes, C.; Kroon, D.; Lepak, N.; Mitchell, S. E.; Peterson, B.; Pressoir, G.; Romero, S.; Rosas, M. O.; Salvo, S.; Yates, H.; Hanson, M.; Jones, E.; Smith, S.; Glaubitz, J. C.; Goodman, M.; Ware, D.; Holland, J. B.; Buckler, E. S. Genetic Properties of the Maize Nested Association Mapping Population. Science 2009, 325 (5941), 737–740. 10.1126/science.1174320.

(41) Lee, M.; Sharopova, N.; Beavis, W. D.; Grant, D.; Katt, M.; Blair, D.; Hallauer, A. Expanding the Genetic Map of Maize with the Intermated B73 × Mo17 (IBM) Population. Plant Mol Biol 2002, 48 (5), 453–461. 10.1023/A:1014893521186.

(42) Hanway, J. J. Growth Stages of Corn (Zea Mays, L.)1. Agronomy Journal 1963, 55 (5), 487–492. 10.2134/agronj1963.00021962005500050024x.

(43) Eichorst, S. A.; Strasser, F.; Woyke, T.; Schintlmeister, A.; Wagner, M.; Woebken, D. Advancements in the Application of NanoSIMS and Raman Microspectroscopy to Investigate the Activity of Microbial Cells in Soils. FEMS Microbiology Ecology 2015, 91 (10), fiv106. 10.1093/femsec/fiv106.

(44) Deng, L.; Fiskal, A.; Han, X.; Dubois, N.; Bernasconi, S. M.; Lever, M. A. Improving the Accuracy of Flow Cytometric Quantification of Microbial Populations in Sediments: Importance of Cell Staining Procedures. Frontiers in Microbiology 2019, 10 (April), 1–13. 10.3389/fmicb.2019.00720.

(45) Bakken, L. R.; Lindahl, V. Recovery of Bacterial Cells from Soil. In Nucleic Acids in the Environment; Trevors, J. T., van Elsas, J. D., Eds.; Springer: Berlin, Heidelberg, 1995; pp 9–27. 10.1007/978-3-642-79050-8_2.

(46) Hu, X.; Zucherman, K. S. Removing Monolayer Cells from Culture Dishes by Incubation with EDTA in Studies of Cell Surface Receptors. Biotechniques 1996, 21 (5), 784, 786. 10.2144/96215bm06.

(47) Günther, S.; Hübschmann, T.; Rudolf, M.; Eschenhagen, M.; Röske, I.; Harms, H.; Müller, S. Fixation Procedures for Flow Cytometric Analysis of Environmental Bacteria. Journal of Microbiological Methods 2008, 75 (1), 127–134. 10/bq9dsz.

(48) Cao, J.-S.; Xu, R.-Z.; Luo, J.-Y.; Feng, Q.; Fang, F. Rapid Quantification of Intracellular Polyhydroxyalkanoates via Fluorescence Techniques: A Critical Review. Bioresource Technology 2022, 350, 126906. 10/gstqvm.

(49) Kawaharasaki, M.; Manome, A.; Kanagawa, T.; Nakamura, K. Flow Cytometric Sorting and RFLP Analysis of Phosphate Accumulating Bacteria in an Enhanced Biological Phosphorus Removal System. Water Science and Technology 2002, 46 (1–2), 139–144. 10.2166/wst.2002.0469.

(50) Zuriani, R.; Vigneswari, S.; Azizan, M. N. M.; Majid, M. I. A.; Amirul, A. A. A High Throughput Nile Red Fluorescence Method for Rapid Quantification of Intracellular Bacterial Polyhydroxyalkanoates. Biotechnology and Bioprocess Engineering 2013, 18 (3), 472–478. 10.1007/s12257-012-0607-z.

(51) Bressan, M.; Trinsoutrot Gattin, I.; Desaire, S.; Castel, L.; Gangneux, C.; Laval, K. A Rapid Flow Cytometry Method to Assess Bacterial Abundance in Agricultural Soil. Applied Soil Ecology 2015, 88, 60–68. 10.1016/j.apsoil.2014.12.007.

(52) Frossard, A.; Hammes, F.; Gessner, M. O. Flow Cytometric Assessment of Bacterial Abundance in Soils, Sediments and Sludge. Front. Microbiol. 2016, 7. 10.3389/fmicb.2016.00903.

(53) Sutermaster, B. A.; Darling, E. M. Considerations for High-Yield, High-Throughput Cell Enrichment: Fluorescence versus Magnetic Sorting. Sci Rep 2019, 9 (1), 227. 10.1038/s41598-018-36698-1.

(54) Ansari, F. A.; Jafri, H.; Ahmad, I.; Abulreesh, H. H. Factors Affecting Biofilm Formation in in Vitro and in the Rhizosphere. In Biofilms in Plant and Soil Health; John Wiley & Sons, Ltd, 2017; pp 275–290. 10.1002/9781119246329.ch15.

(55) Büks, F.; Kaupenjohann, M. Enzymatic Biofilm Digestion in Soil Aggregates Facilitates the Release of Particulate Organic Matter by Sonication. SOIL 2016, 2 (4), 499–509. 10.5194/soil-2-499-2016.

(56) Metz, S.; Dos Santos, A. L.; Berman, M. C.; Bigeard, E.; Licursi, M.; Not, F.; Lara, E.; Unrein, F.; Lopes dos Santos, A.; Berman, M. C.; Bigeard, E.; Dos Santos, A. L.; Berman, M. C.; Bigeard, E.; Licursi, M.; Not, F.; Lara, E.; Unrein, F. Diversity of Photosynthetic Picoeukaryotes in Eutrophic Shallow Lakes as Assessed by Combining Flow Cytometry Cell-Sorting and High Throughput Sequencing. FEMS Microbiology Ecology 2019, 95 (5), 1–12. 10.1093/femsec/fiz038.

(57) Daims, H. Use of Fluorescence in Situ Hybridization and the Daime Image Analysis Program for the Cultivation-Independent Quantification of Microorganisms in Environmental and Medical Samples. Cold Spring Harbor Protocols 2009, 4 (7), 1–8. 10.1101/pdb.prot5253.

(58) He, P.; Son, Y.; Berkowitz, J.; Li, G.; Lee, J.; Han, I.; Craft, E.; Piñeros, M.; Kao-Kniffin, J.; Gu, A. Z. Recycled Phosphorus Bioamendments from Wastewater Impact Rhizomicrobiome and Benefit Crop Growth: Sustainability Implications at Water-Food Nexus. Environ. Sci. Technol. 2025, 59 (4), 2131–2143. 10.1021/acs.est.4c07901.

(59) Li, Y.; Rahman, S. M.; Li, G.; Fowle, W.; Nielsen, P. H.; Gu, A. Z. The Composition and Implications of Polyphosphate-Metal in Enhanced Biological Phosphorus Removal Systems. Environ. Sci. Technol. 2019, 53 (3), 1536–1544. 10.1021/acs.est.8b06827.

(60) Wang, D.; Li, Y.; Cope, H. A.; Li, X.; He, P.; Liu, C.; Li, G.; Rahman, S. M.; Tooker, N. B.; Bott, C. B.; Onnis-Hayden, A.; Singh, J.; Elfick, A.; Marques, R.; Jessen, H. J.; Oehmen, A.; Gu, A. Z. Intracellular Polyphosphate Length Characterization in Polyphosphate Accumulating Microorganisms (PAOs): Implications in PAO Phenotypic Diversity and Enhanced Biological Phosphorus Removal Performance. Water Research 2021, 206, 117726. 10.1016/j.watres.2021.117726.

(61) Chavan, S.; Yadav, B.; Tyagi, R. D.; Drogui, P. A Review on Production of Polyhydroxyalkanoate (PHA) Biopolyesters by Thermophilic Microbes Using Waste Feedstocks. Bioresource Technology 2021, 341, 125900. 10/gr8kr7.

(62) Chang, C. F.; Shuman, H.; Somlyo, A. P. Electron Probe Analysis, X-Ray Mapping, and Electron Energy-Loss Spectroscopy of Calcium, Magnesium, and Monovalent Ions in Log-Phase and in Dividing Escherichia Coli B Cells. J Bacteriol 1986, 167 (3), 935–939. 10.1128/jb.167.3.935-939.1986.

(63) Smith, R. J. Calcium and Bacteria. Adv Microb Physiol 1995, 37, 83–133. 10.1016/s0065-2911(08)60144-7.

(64) Barra Caracciolo, A.; Grenni, P.; Cupo, C.; Rossetti, S. In Situ Analysis of Native Microbial Communities in Complex Samples with High Particulate Loads. FEMS Microbiology Letters 2005, 253 (1), 55–58. 10.1016/j.femsle.2005.09.018.

(65) Kamika, I.; Coetzee, M.; Mamba, B. B.; Msagati, T.; Momba, M. N. B. The Impact of Microbial Ecology and Chemical Profile on the Enhanced Biological Phosphorus Removal (EBPR) Process: A Case Study of Northern Wastewater Treatment Works, Johannesburg. International Journal of Environmental Research and Public Health 2014, 11 (3), 2876–2898. 10.3390/ijerph110302876.

(66) Nielsen, P. H.; McIlroy, S. J.; Albertsen, M.; Nierychlo, M. Re-Evaluating the Microbiology of the Enhanced Biological Phosphorus Removal Process. Current Opinion in Biotechnology 2019, 57 (Figure 1), 111–118. 10.1016/j.copbio.2019.03.008.

(67) Stokholm-Bjerregaard, M.; McIlroy, S. J.; Nierychlo, M.; Karst, S. M.; Albertsen, M.; Nielsen, P. H. A Critical Assessment of the Microorganisms Proposed to Be Important to Enhanced Biological Phosphorus Removal in Full-Scale Wastewater Treatment Systems. Frontiers in Microbiology 2017, 8 (APR), 1–18. 10.3389/fmicb.2017.00718.

(68) Bai, Y.; Zhang, Y.; Quan, X.; Chen, S. Nutrient Removal Performance and Microbial Characteristics of a Full-Scale IFAS-EBPR Process Treating Municipal Wastewater. Water Science and Technology 2016, 73 (6), 1261–1268. 10.2166/wst.2015.604.

(69) Singleton, C. M.; Petriglieri, F.; Wasmund, K.; Nierychlo, M.; Kondrotaite, Z.; Petersen, J. F.; Peces, M.; Dueholm, M. S.; Wagner, M.; Nielsen, P. H. The Novel Genus, ‘Candidatus Phosphoribacter’, Previously Identified as Tetrasphaera, Is the Dominant Polyphosphate Accumulating Lineage in EBPR Wastewater Treatment Plants Worldwide. ISME Journal 2022, 16 (6), 1605–1616. 10.1038/s41396-022-01212-z.

(70) Pankratov, T. A.; Grouzdev, D. S.; Patutina, E. O.; Kolganova, T. V.; Suzina, N. E.; Berestovskaya, J. J. Lichenibacterium Ramalinae Gen. Nov, Sp. Nov., Lichenibacterium Minor Sp. Nov., the First Endophytic, Beta-Carotene Producing Bacterial Representatives from Lichen Thalli and the Proposal of the New Family Lichenibacteriaceae within the Order Rhizobiales. Antonie van Leeuwenhoek 2020, 113 (4), 477–489. 10/gstz8p.

(71) Wang, H.-G.; Huang, H.; Liu, R.-L.; Mao, Y.-P.; Biswal, B. K.; Chen, G.-H.; Wu, D. Investigation on Polyphosphate Accumulation in the Sulfur Transformation-Centric EBPR (SEBPR) Process for Treatment of High-Temperature Saline Wastewater. Water Research 2019, 167, 115138. 10/gmngmh.

(72) Lima, D. C.; Aviles, A. C.; Alpers, R. T.; Perkins, A.; Schoemaker, D. L.; Costa, M.; Michel, K. J.; Kaeppler, S.; Ertl, D.; Romay, M. C.; Gage, J. L.; Holland, J.; Beissinger, T.; Bohn, M.; Buckler, E.; Edwards, J.; Flint-Garcia, S.; Gore, M. A.; Hirsch, C. N.; Knoll, J. E.; McKay, J.; Minyo, R.; Murray, S. C.; Schnable, J.; Sekhon, R. S.; Singh, M. P.; Sparks, E. E.; Thomison, P.; Thompson, A.; Tuinstra, M.; Wallace, J.; Washburn, J. D.; Weldekidan, T.; Xu, W.; de Leon, N. 2020-2021 Field Seasons of Maize GxE Project within the Genomes to Fields Initiative. BMC Res Notes 2023, 16 (1), 219. 10.1186/s13104-023-06430-y.

(73) Kurashita, H.; Hatamoto, M.; Tomita, S.; Yamaguchi, T.; Narihiro, T.; Kuroda, K. Comparative Metagenomic Analysis Provides Novel Insights into the Ecophysiology of Myxococcota in Activated Sludge Systems. bioRxiv April 28, 2024, p 2024.04.26.591422. 10.1101/2024.04.26.591422.

(74) Zhou, T.; Xiang, Y.; Liu, S.; Ma, H.; Shao, Z.; He, Q.; Chai, H. Microbial Community Dynamics and Metagenomics Reveal the Potential Role of Unconventional Functional Microorganisms in Nitrogen and Phosphorus Removal Biofilm System. Science of The Total Environment 2023, 905, 167194. 10.1016/j.scitotenv.2023.167194.

(75) Akone, S. H.; Hug, J. J.; Kaur, A.; Garcia, R.; Müller, R. Structure Elucidation and Biosynthesis of Nannosterols A and B, Myxobacterial Sterols from Nannocystis Sp. MNa10993. Journal of Natural Products 2023, 86 (4), 915. 10.1021/acs.jnatprod.2c01143.

(76) Peiffer, J. A.; Spor, A.; Koren, O.; Jin, Z.; Tringe, S. G.; Dangl, J. L.; Buckler, E. S.; Ley, R. E. Diversity and Heritability of the Maize Rhizosphere Microbiome under Field Conditions. Proceedings of the National Academy of Sciences 2013, 110 (16), 6548–6553. 10.1073/pnas.1302837110.

(77) Pérez-García, L.-A.; Sáenz-Mata, J.; Fortis-Hernández, M.; Navarro-Muñoz, C. E.; Palacio- Rodríguez, R.; Preciado-Rangel, P. Plant-Growth-Promoting Rhizobacteria Improve Germination and Bioactive Compounds in Cucumber Seedlings. Agronomy 2023, 13 (2), 315. 10.3390/agronomy13020315.

(78) Kumar Arora, N.; Fatima, T.; Mishra, J.; Mishra, I.; Verma, S.; Verma, R.; Verma, M.; Bhattacharya, A.; Verma, P.; Mishra, P.; Bharti, C. Halo-Tolerant Plant Growth Promoting Rhizobacteria for Improving Productivity and Remediation of Saline Soils. Journal of Advanced Research 2020, 26, 69–82. 10.1016/j.jare.2020.07.003.

(79) Dai, W.; Liu, M.; Wang, N.; Ye, X.; Liu, Y.; Yao, D.; Wang, L.; Cui, Z.; Yan, P.; Cheng, C.; Huang, Z.; Wang, H. Positive Contribution of Predatory Bacterial Community to Multiple Nutrient Cycling and Microbial Network Complexity in Arsenic-Contaminated Soils. Applied Soil Ecology 2023, 185, 104792. 10.1016/j.apsoil.2022.104792.

(80) Yang, X.; Gao, R.; Zhang, Q.; Yung, C. C. M.; Yin, H.; Li, J. Quantification of Polyphosphate in Environmental Planktonic Samples Using a Novel Fluorescence Dye JC- D7. Environ. Sci. Technol. 2024, 58 (32), 14249–14259. 10.1021/acs.est.4c04545.

