## Supplementary Information for "Discovering Polyphosphate and Polyhydroxyalkanoate-Accumulating Organisms Across Ecosystems: Phenotype-targeted Genotyping via FACS-sequencing"

### Text S1 Activation of sludge samples from enhanced biological phosphorus removal (EBPR)

Sludge samples were activated using phosphorus (P) release and uptake tests in accordance with Gu et al., 2008 using acetate as a carbon source.^1^ Specifically, activated sludge from the aeration tank was transferred to a batch reactor and supplemented with sodium acetate to a concentration of 80 mg L^−1^. Anaerobic conditions (DO < 0.01 mg L^−1^) were created for 120 min by purging the sludge with N_2_ gas, and the activity of polyphosphate (polyP)-accumulating organisms (PAOs) related to P-release and carbon uptake was measured. Subsequently, aerobic conditions (DO > 2.0 mg L^−1^) were established for 180 min by purging the sludge with air. Samples were collected for soluble chemical oxygen demand (sCOD) and PO_4_^-^-P measurements. The P release and uptake profiles for three EBPR plants are shown in Figure S2.

### Text S2 FACS gating strategy for PHA-PAO targeted single-cell sorting

To ensure only cells are identified and sorted at the point of interrogation during FACS, we performed two more steps. Firstly, leveraging the forward scattering channel (FSC) hardware of FACS to ensure targeted sorting of single-cells and discriminating against remaining aggregates, we constrained acceptable particle size by forward scattering height (FSC-H) and forward scattering area (FSC-A) ratios based on pure culture single-cell suspensions of known PAO, *Phycicoccus elongatus* (previously named *Tetrasphaera elongata,* Lp2, DSM No. 14184) (Figure S5, step 1). Following cell-sized event identification, background noise and autofluorescence was removed by comparing event count histograms from unstained negative controls and single-stained SYBR Green I (SG) positive controls in the SG fluorescent channel, excluding 99.5% of all events within the unstained negative control (Figure S5, step 2).

Secondly, we implemented a sequential gating strategy to target cells exhibiting a characteristic triple-positive fluorescent signal, using single- and dual-stained positive controls to set fluorescence thresholds for identifying and sorting PHA-PAOs. Gating boundaries were first established with a single-stained SG-positive control to determine the threshold for NR-negative samples and account for any spectral spillover (Figure S5 step 3, 4). This approach was then applied to polyP-containing cells using SG+NR dual-stained samples to set TC-negative thresholds (Figure S5, step 5). Finally, candidate PHA-PAOs containing both intracellular polyP and PHA were identified by selecting cells that were both NR-positive and TC-positive, as indicated in Figure S5 (Step 5, blue).

### Text S3 16S rRNA Gene Amplicon Sequencing

Each primer contained a unique six-base index sequence for sample multiplexing, along with Illumina flow cell binding and sequencing sites.^4^ The 25 μL PCR mix included 1× ThermoPol buffer, 0.2 μM forward primer, 0.2 μM reverse primer, 200 μM total dNTPs, 15 μg bovine serum albumin, 0.625 units *Taq* DNA polymerase (New England Biolabs, USA), and 10 ng of template DNA. 25 μL PCR reactions were performed in triplicate as follows: 95°C for 3 min, 35 cycles of 95°C for 30 sec, 50°C for 30 sec, 68°C for 1 min, and a final extension of 68°C for 7 min. Equal quantities of 16S rRNA gene amplicons were pooled, excised from an agarose gel, and purified using the Wizard SV Gel and PCR Clean-Up System (Promega, WI, USA). A 3 pM library containing 10% PhiX Control v3 (Illumina, CA, USA) was sequenced on a MiSeq (Illumina, CA, USA) using a 2 x 250 cycle MiSeq Reagent Kit v2.

The paired-end reads were processed using two distinct pipelines tailored to sample type. For sludge samples, sequences were assembled, merged, and clustered into operational taxonomic units (OTUs) at 97% similarity using Mothur. Representative sequences were classified against the SILVA database.^5^

Soil sample sequences were processed using the DADA2 pipeline. Reads were filtered with a maximum expected error rate (maxEE) of 2, and the last base of each read was trimmed to mitigate low-quality regions. Amplicon sequence variants (ASVs) were inferred, and sequences were retained if they were between 252–254 bp in length. Chimeric, chloroplast, and mitochondrial sequences were removed.^6,7^ Samples with fewer than 1,000 reads were excluded from downstream analyses. To account for varying sequencing depths, rarefaction was performed using iNEXT prior to alpha diversity assessments ().^8^

### Text S4 Classification of Detected Genera by Polyphosphate and PHA Accumulation Potential

A hierarchical color-coding system was established to categorize genera based on their documented PAO-related functionality:

Red Stars (★): Genera with experimentally confirmed anaerobic phosphorus release and aerobic phosphorus uptake characteristics as reported in peer-reviewed literature. These organisms demonstrated classic PAO behavior through controlled laboratory studies or field observations.

Yellow Stars (★): Genera with documented experimental evidence of polyphosphate (polyP) and polyhydroxyalkanoate (PHA) accumulation but without conclusive demonstration of PAO activity. These classifications were based on microscopic observations, staining techniques, or biochemical assays showing intracellular polymer storage.

Green Stars (★): Genera identified as having genetic potential for PHA-PAO metabolism through genomic analysis. Representative genomes were retrieved from public genome databases including the National Center for Biotechnology Information (NCBI) GenBank and BioCyc Database Collection. Genomes were computationally screened for the presence of target functional genes associated with polyphosphate kinase (ppk), exopolyphosphatase (ppx), PHA synthase (phaC), and other key enzymes involved in PAO metabolism pathways.

Blue Stars (★): For taxonomic assignments that could not be resolved to the genus level due to sequencing limitations, classifications were made at higher taxonomic ranks (family or order level). In these cases, all genera within the respective family or order were systematically reviewed for reported polyP and PHA accumulation capabilities. However, these assignments received blue star designations to indicate the indirect nature of the functional inference.

### Test S5 Single cell Raman spectroscopy (SCRS)

Bacteria were extracted and separated from greenhouse maize pot samples using density-gradient separation according to a previously developed protocol.^9^ The collected bacterial solution was diluted appropriately, and 1 μL was added onto CaF2 Raman slides (Crystran Ltd., UK). Raman spectra were acquired using a Horiba LabRam Evolution (Horiba, Kyoto, Japan) equipped with a 50× objective (Olympus LMPLFLN 50×, Tokyo, Japan) following established protocols.^10–12^

Microbial Raman spectra were distinguished from abiotic particles using biological signature peaks at 1003 and 1657 cm⁻¹, corresponding to phenylalanine and amide I, respectively.^12^ At least 100 microbial Raman spectra were acquired per sample based on SCRS sample size sufficiency analysis.^10,11^ PAO cells that contain polyphosphate were identified via the signature peaks at 690 and 1170 cm^–1^ according to our previous studies for PAO identification.^13^ The relative abundance of PAOs was calculated by dividing the number of PAO-identified spectra by the total spectra acquired per sample.

### Test S6 Preliminary Evaluation of PolyP-Containing Organisms (PCOs) in Soil Samples

Prior to implementing the TriFlow protocol in soil samples, we initially validated polyP-DNA dual staining (without the use of NR) by comparing the relative abundance of PCOs identified through fluorescence-activated cell sorting (FACS) with those determined by SCRS in a separate set of rhizosphere soil samples. A significant positive monotonic relationship was observed between the two techniques (Spearman’s *ρ* = 0.61, *p* = 0.002), demonstrating statistically robust concordance in detecting PCOs (Figure S7). The data presented here constitute a subset of samples from a manuscript currently under preparation. Detailed descriptions are provided in that document and are not repeated here to maintain focus in the main text.

The rhizosphere soil samples were collected during the summer of 2021 from farms in upstate New York, United States. The farms and their respective crop type were as follows: 1). Farm 1 (1443 Ridge Rd, Penn Yan, NY 14527; 42.6929190566136° N, 76.99673010440144° W) – rhizosphere soils from organically grown corn, wheat, soybean, alfalfa, and triticale; 2). Farm 2 (780 Ridge Rd, Penn Yan, NY 14527; 42.72775159642192° N, 77.00261800255171° W) – rhizosphere soils from conventionally grown corn, wheat, soybean, alfalfa, and triticale; 3). Farm 3 (3320 NY-215, Cortland, NY 13045; 42.568046701646296° N, 76.19280911605144° W) – rhizosphere soils from organically grown potato, tomato, carrot, pea, and oat; 4). Farm 4 (1157 County Rd 39, Bainbridge, NY 13733; 42.28082210297718° N, 75.47726548907552° W) - rhizosphere soils from conventionally grown potato, tomato, and carrot; 5). Farm 5 (1798 Co Rd 4, Seneca Castle, NY 14547; 42.88706285558538° N, 77.07643635889123° W) – rhizosphere soils from organically grown lettuce and conventionally grown squash.

For each crop mentioned above, five seedlings were destructively sampled, and the rhizosphere soil adhering to their roots was carefully collected. All rhizosphere soil samples were transported in a pre-chilled cooler with ice packs to maintain temperature. Upon arrival at the lab, samples were immediately processed for FACS. Due to the labor- and time-intensive nature of FACS, a single composite sample, pooled from the five replicates, was sorted for each crop and farm type. However, flow cytometric couting analysis was performed on all five replicates to minimize potential bias. Samples were stained with SG for DNA and TC for polyP, while Nile Red was not used in this batch of samples due to differing experimental purposes.

SCRS, which targets the molecular composition of microbial cells, has demonstrated exceptional sensitivity and accuracy in detecting and quantifying intracellular polymers, such as polyP, PHA, and glycogen.^13^ This capability allows for effective quantification and characterization of PAO communities.^14,15^ The alignment of PAO percentages determined by FACS with those identified through an independent technique further underscores the reliability of FACS in quantifying PAO communities within complex environmental matrices.

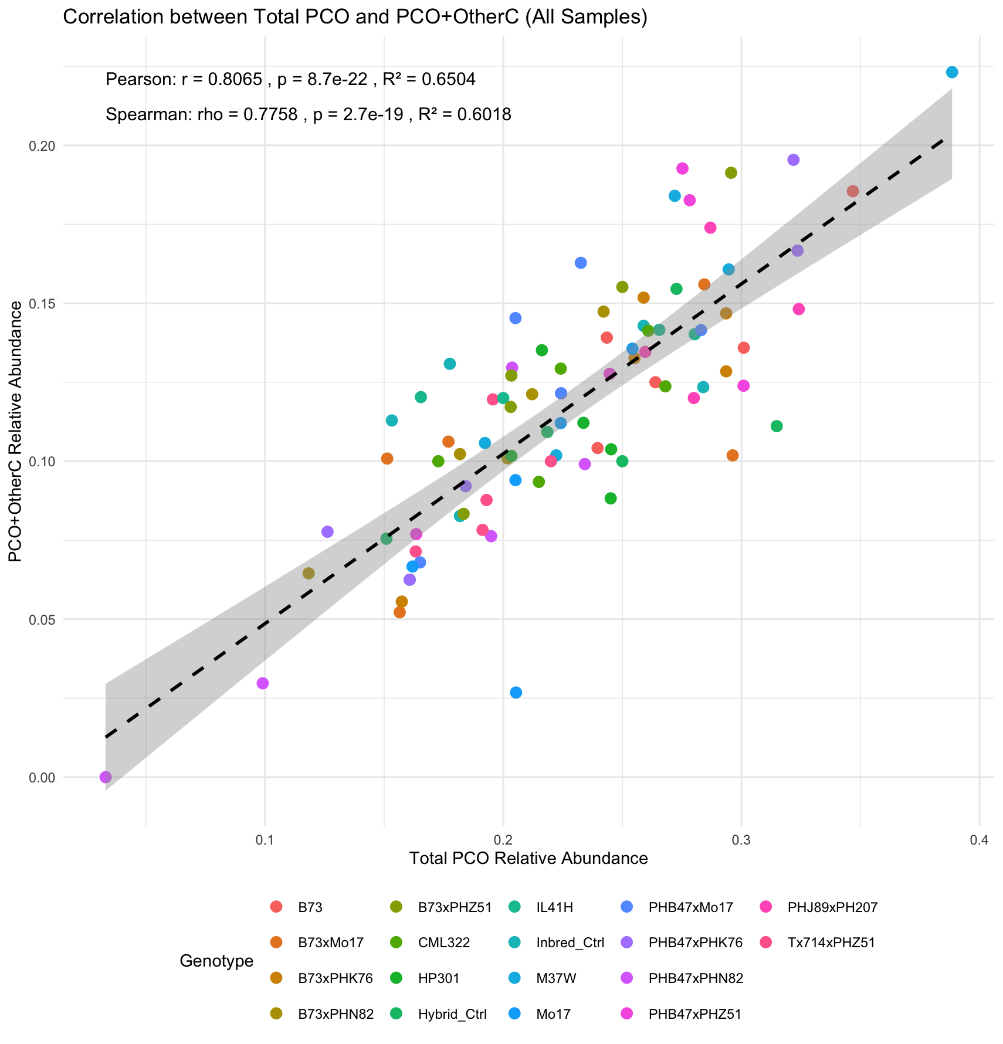

Figure S1 Strong correlation between the abundance of polyphosphate (polyP)-containing organisms (PCOs) and polyP-accumulating organisms (PAOs) in maize rhizosphere, as detected by single-cell Raman spectroscopy. Note: As defined in the main text, PCOs are organisms accumulating polyP irrespective of intracellular carbon polymer content, while PAOs co-accumulate polyP and intracellular C, primarily PHA, with surplus capacity. Sampling methodology is detailed in Text S4 and Table S1.

A)

B)

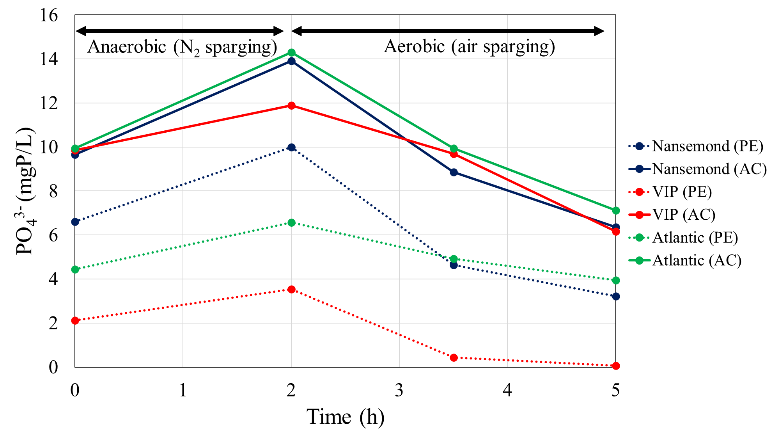

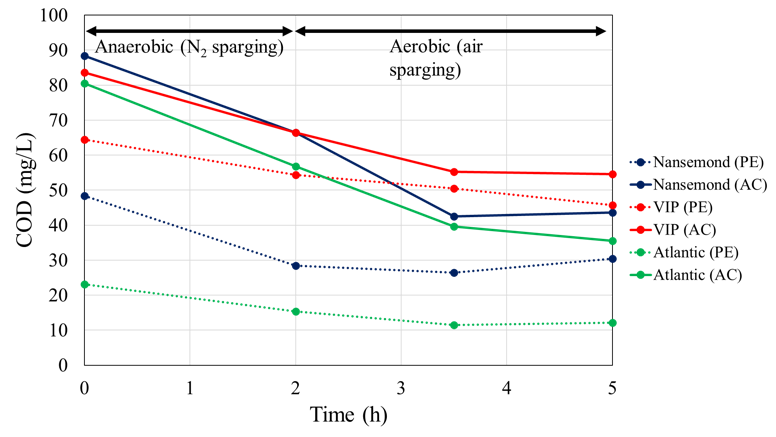

Figure S2 Profiles of (A) orthophosphate and (B) chemical oxygen demand (COD) during cyclic anaerobic/aerobic batch tests using returned activated sludge (RAS) sampled from three full-scale enhanced biological phosphorus removal (EBPR) plants (Nansemond, VIP and Atlantic).

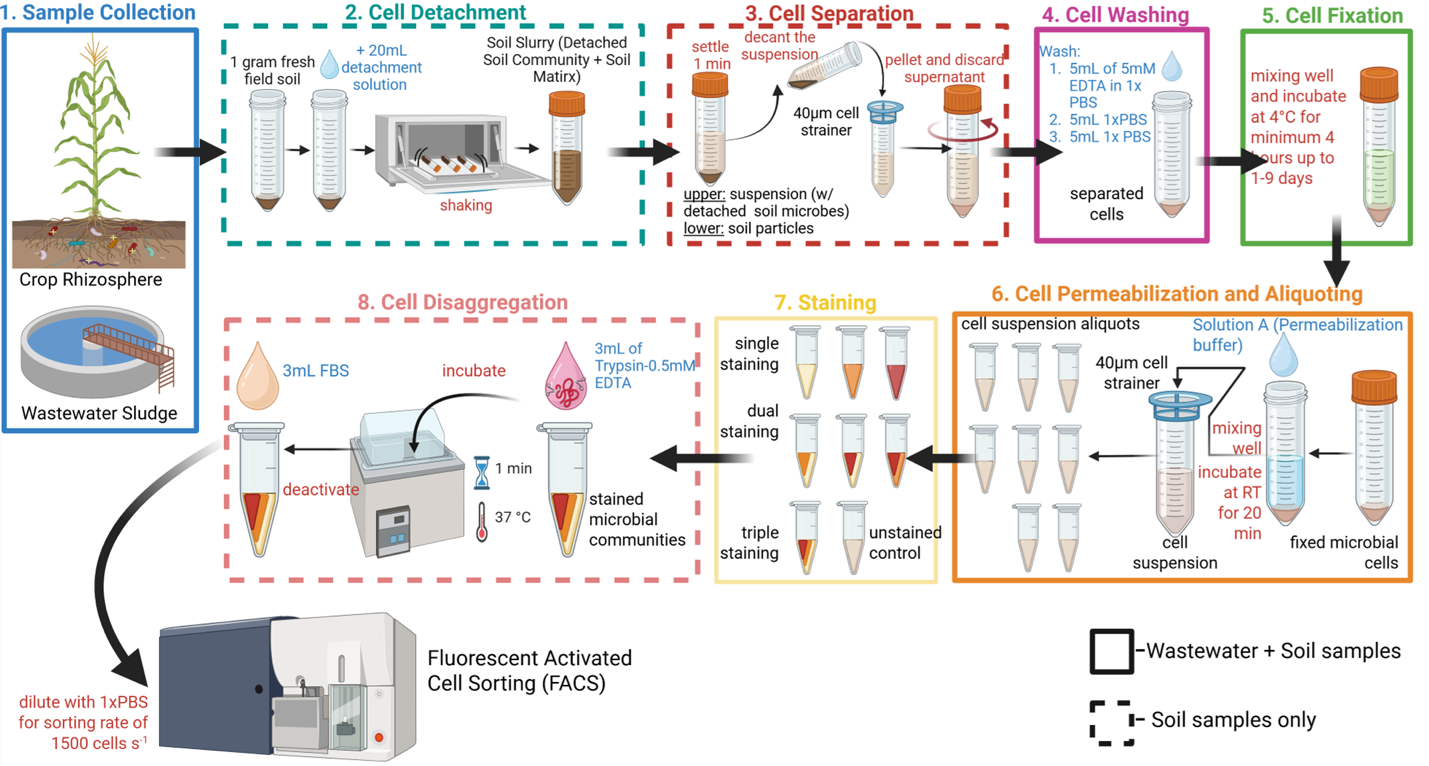

Figure S3 Overview of the experimental workflow for microbial cell isolation, pre-treatment, and staining for fluorescent activated cell sorting (FACS) from crop rhizosphere soils and wastewater sludge. Steps specific to rhizosphere soil samples are indicated with dashed borders, while steps common to both sample types are shown with solid borders. Text in red indicates an action being undertaken, text in blue indicates the solution being used, and text in black are descriptions.

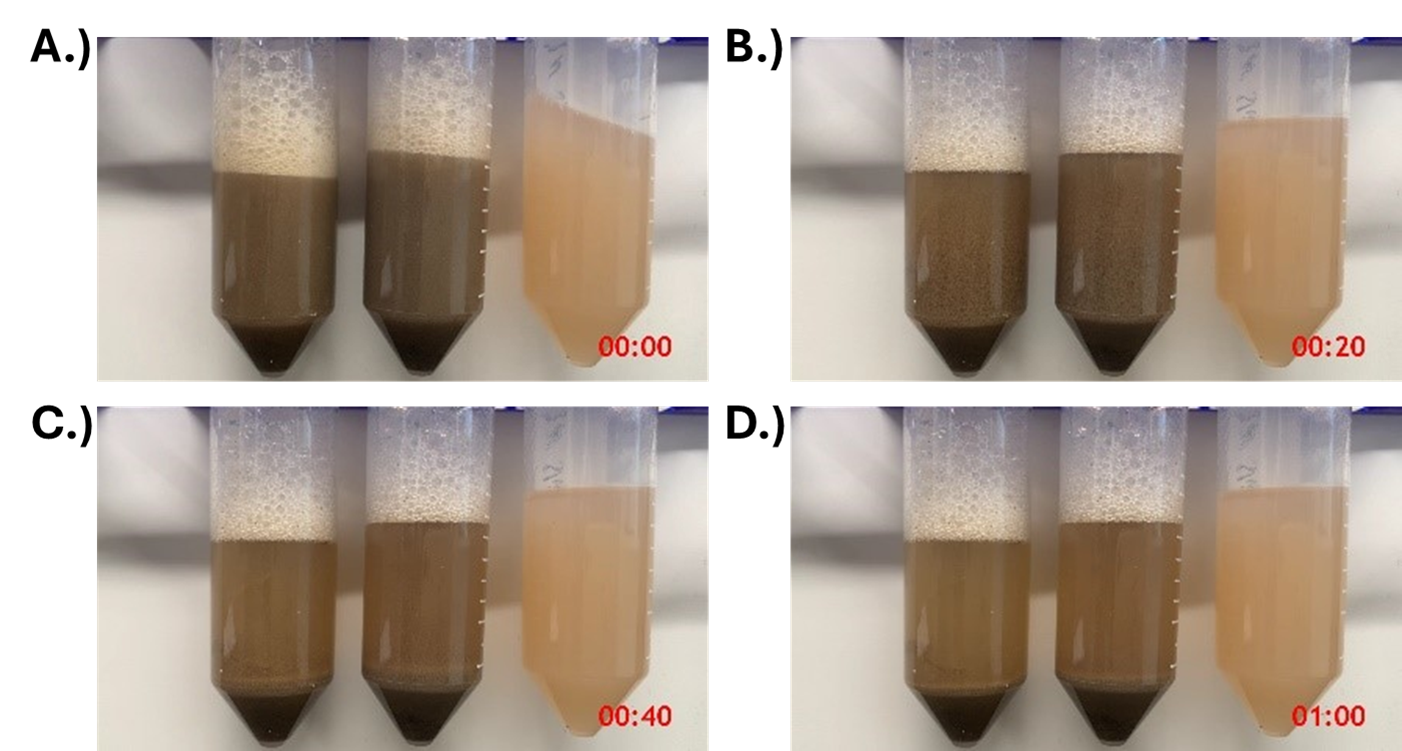

Figure S4 Frame by frame analysis of sedimentation time optimization of soil-cell separation following detergent-based cell detachment comparing settling of: from left to right, two soil replicates and mixed culture from high strength synthetic wastewater fed bioreactor. Time stamps in red at bottom right corner of each frame

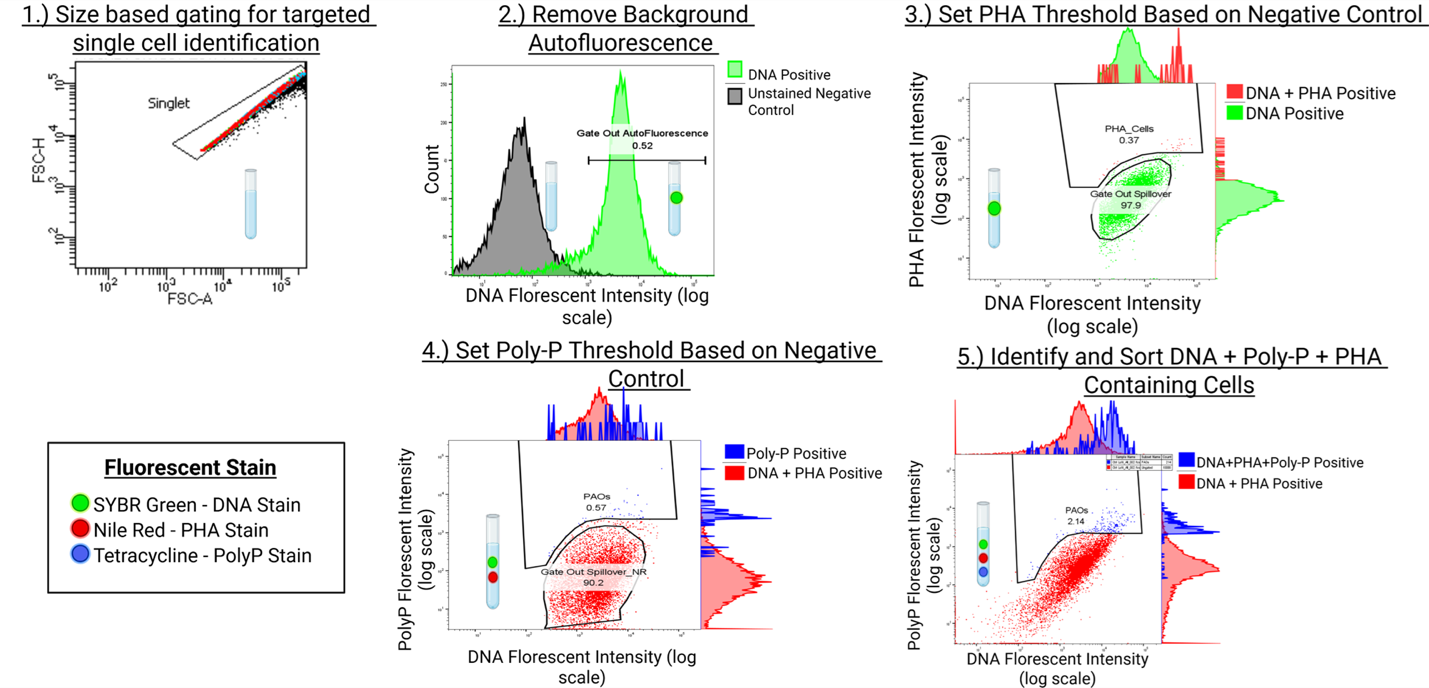

Figure S5 Example gating panel for identification and quantification of single-cell candidate triple-stained positive cells using TriFlow-Seq.

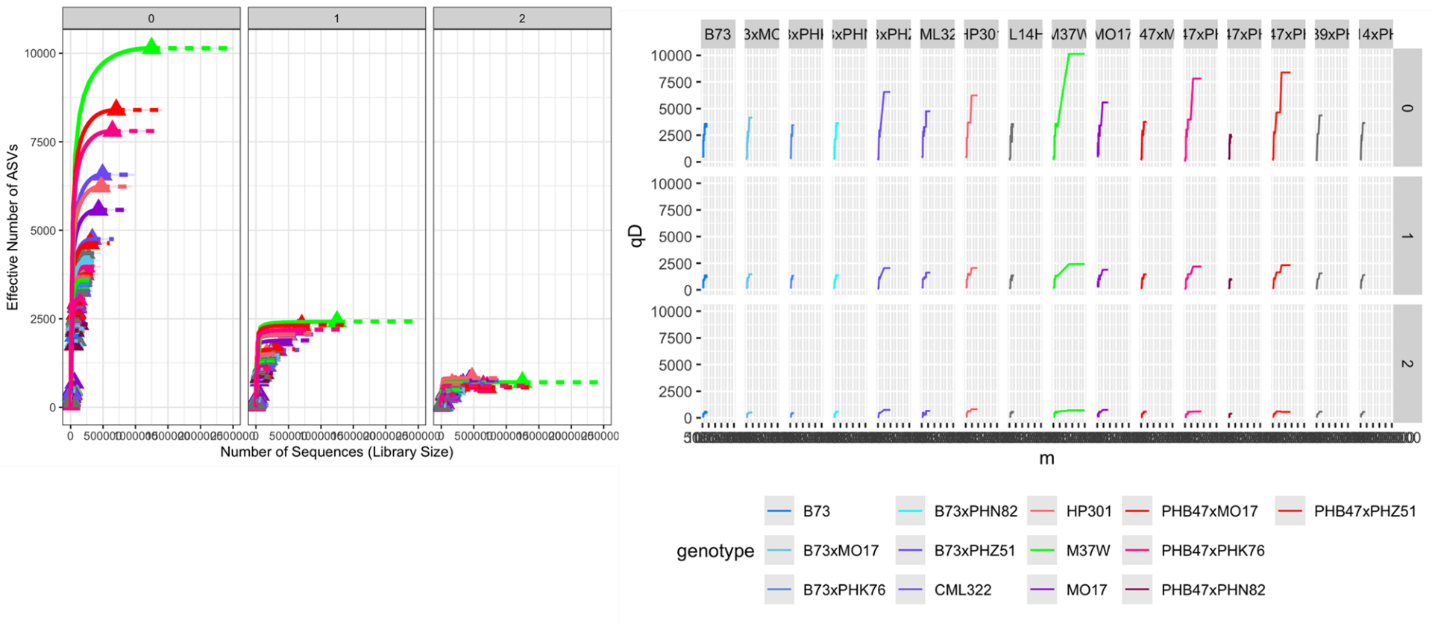

Figure S6 Rarefaction curves generated by iNEXT for sorted rhizomicrobiome samples, demonstrating that TriFlow-Seq achieved adequate sequencing depth for community analysis

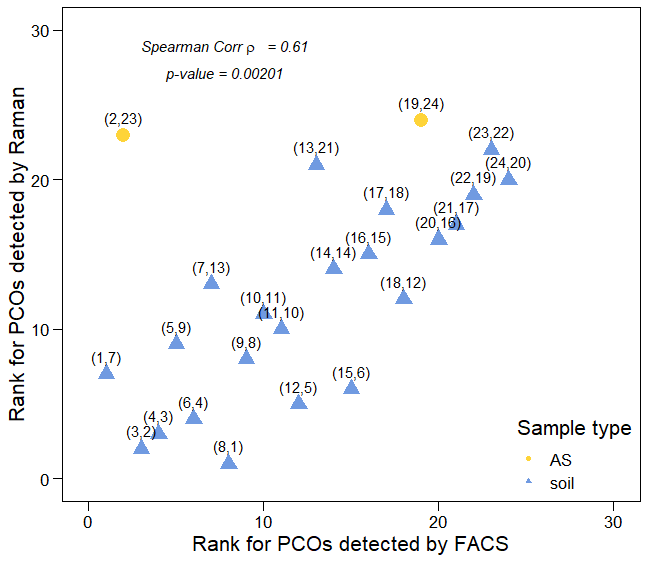

Figure S7 Spearman's rank correlation coefficients of polyphosphate (polyp)-containing organisms (PCOs) detected by fluorescence-activated cell sorting (FACS) and Single-Cell Raman Spectroscopy (SCRS), irrespective of PHA presence. The strong correlation between the two independent techniques validates the effectiveness of TriFlow in detecting and quantifying PCOs. Sample information is listed in Text S4.

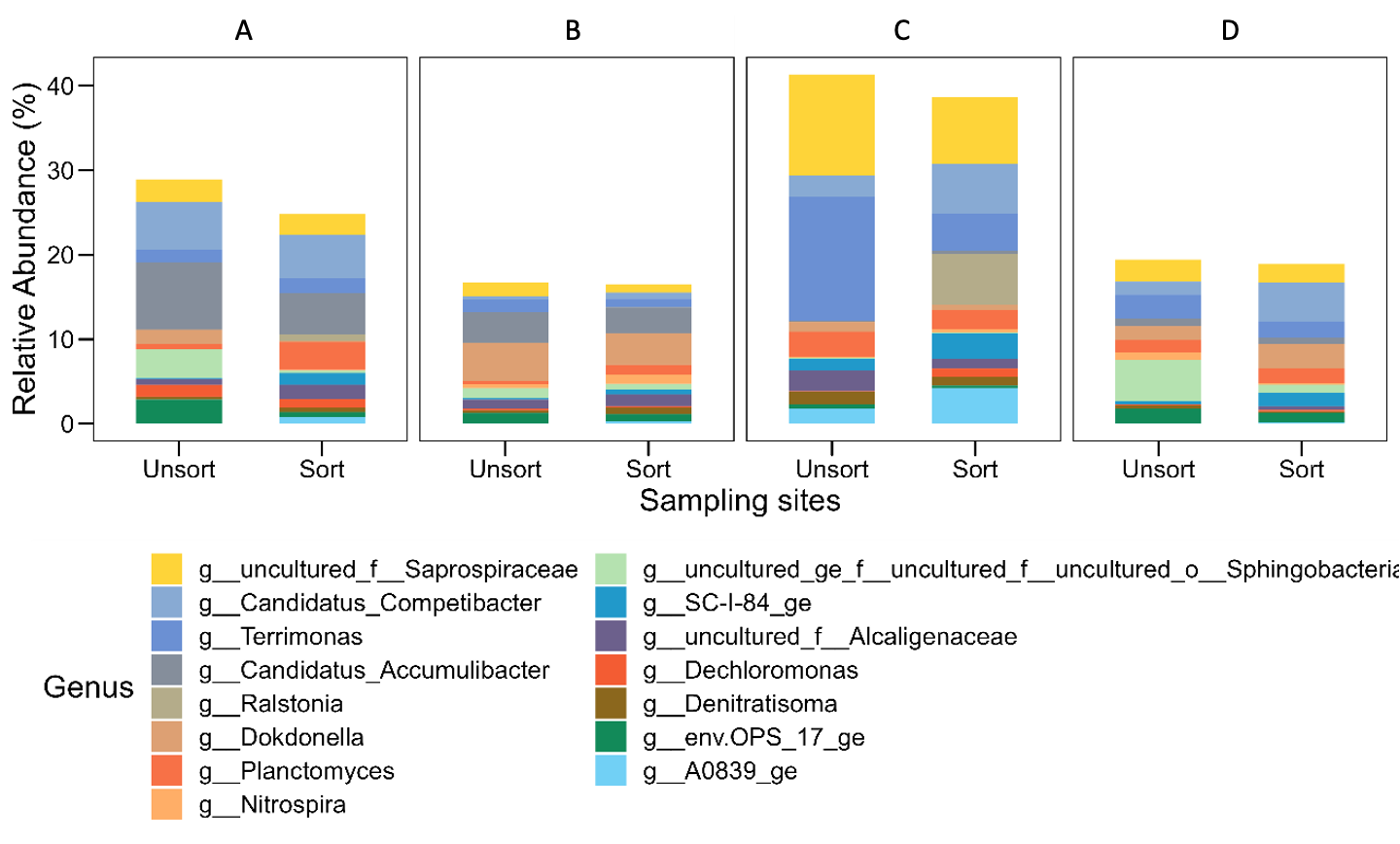

Figure S8 Relative abundances of the microbial communities (genus level) in three full-cale EBPR plants and one pilot-scale S2EBPR plant in the U.S. before and after FACS sorting. These samples were activated with P release and uptake tests. Relative abundances below 0.1% are not shown.

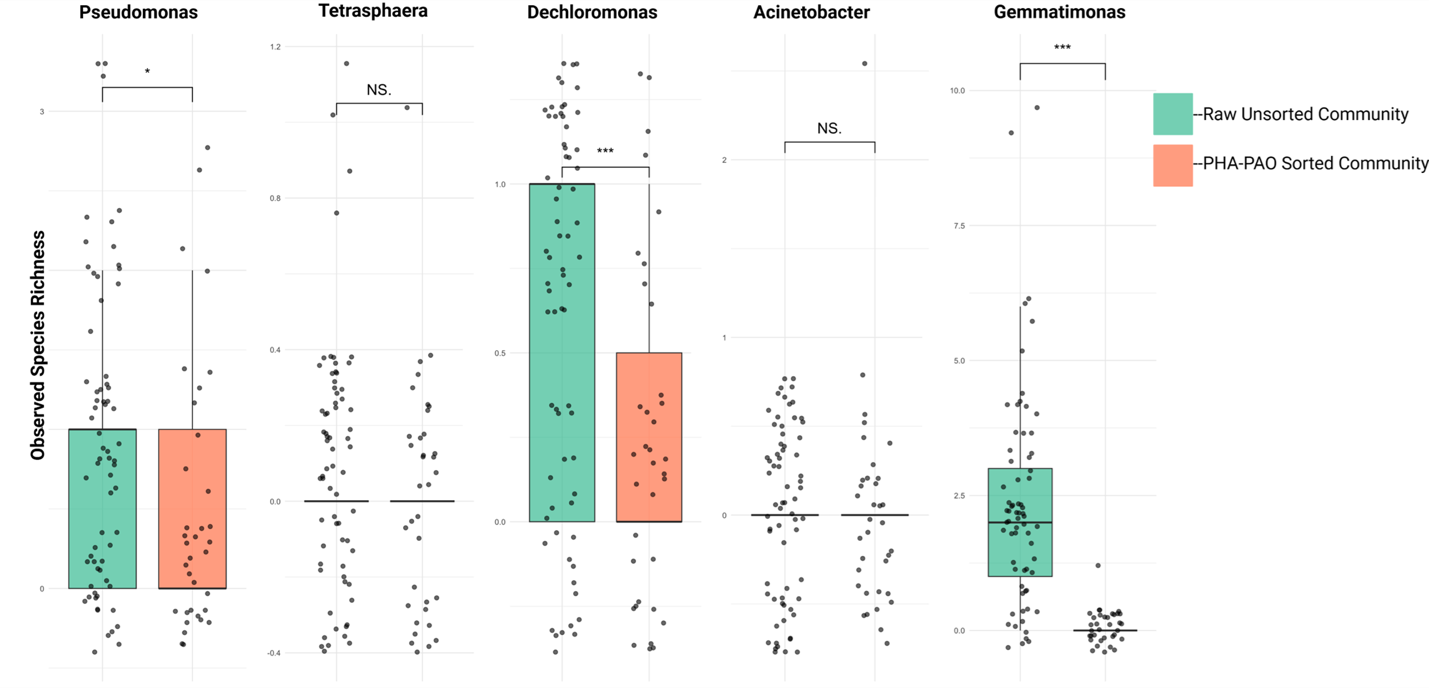
 Figure S9 Unrarefied, observed species richness of canonical PAOs detected across raw unsorted maizr rhizosphere (green) and PHA-PAO sorted rhizomicrobiome (orange). Signficnace based on t.test with a *p* <0.05 indicated by *, *p* <0.01 by **, and *p* <0.001 by ***.

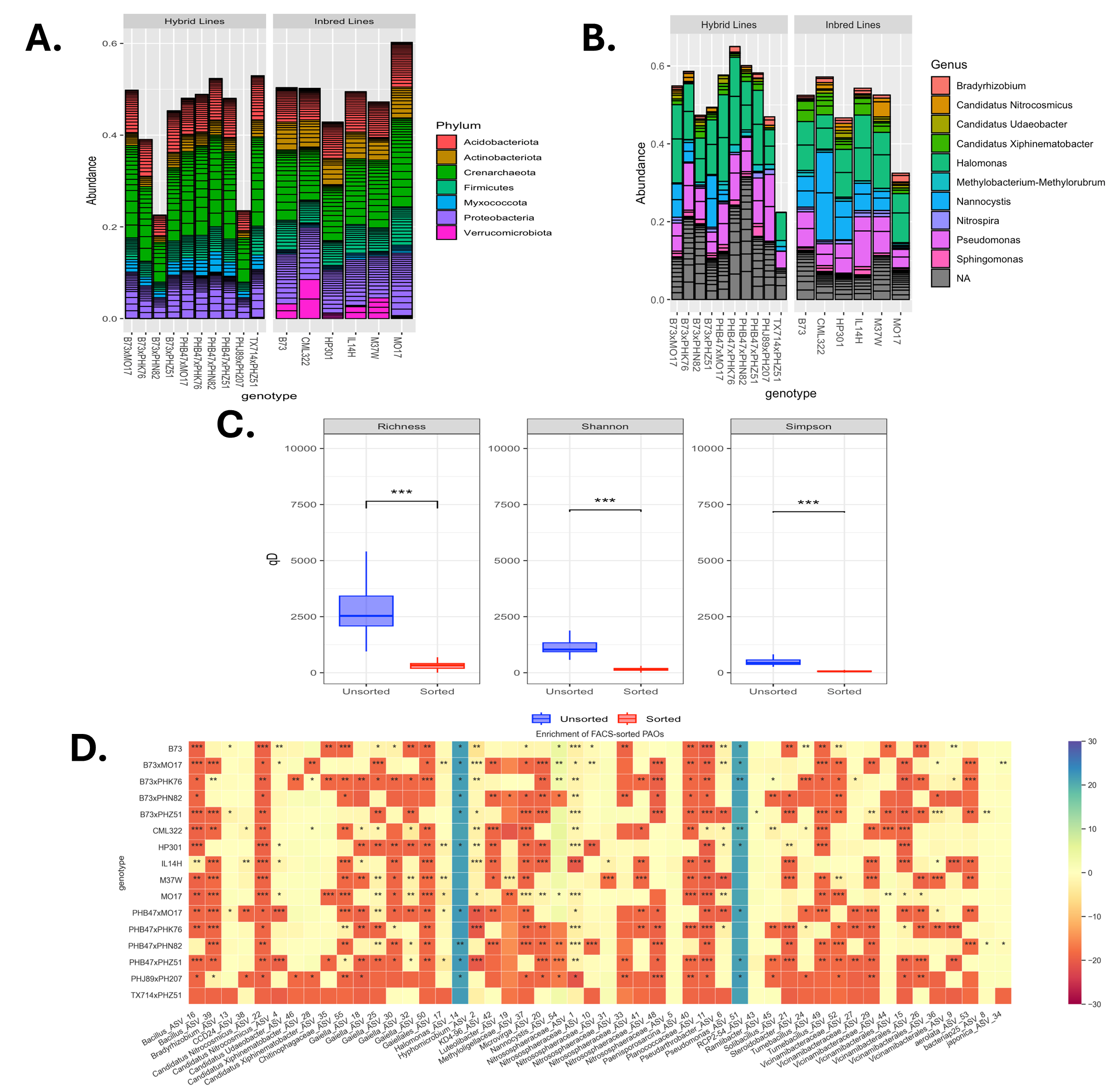

Figure S10 Observed Hill numbers (from left to right: species richness, exponential of Shannon index, and inverse of Simpson index) for rarefied TriFlow-seq-identified polyphosphate (polyP)-accumulating organisms (PAOs) and unsorted rhizomicrobiome communities, estimated using iNEXT.

Boxplots display the median diversity (central line), interquartile range (box), and minimum/maximum values (whiskers). Statistical significance between groups was determined using the Kruskal-Wallis test, with *** denoting a *p*-value <0.001.

Table S1. Rhizosphere soil smapled from maize inbred and hybrid lines.

| Soil Series | Maize Genotypes | Total plants collected |
| --- | --- | --- |
| Lima silt soam (0-3% slope) | B73 | 4 |
|  | Mo17 | 4 |
|  | HP301 | 4 |
|  | IL14H | 4 |
|  | CML322 | 4 |
|  | M37W | 4 |
| Honeoye silt loam (3-8% slope) and Kendaia & Lyons silt loam (0-3% slope) | B73xMo17 | 6 |
|  | TX714xPHZ51 | 6 |
|  | PHB47xPHK76 | 6 |
|  | PHB47xPHZ51 | 6 |
|  | B73xPHZ51 | 6 |
|  | PHB47xMo17 | 6 |
|  | PHJ89xPH207 | 6 |
|  | B73xPHN82 | 6 |
|  | PHB47xPHN82 | 6 |
|  | B73xPHK76 | 6 |

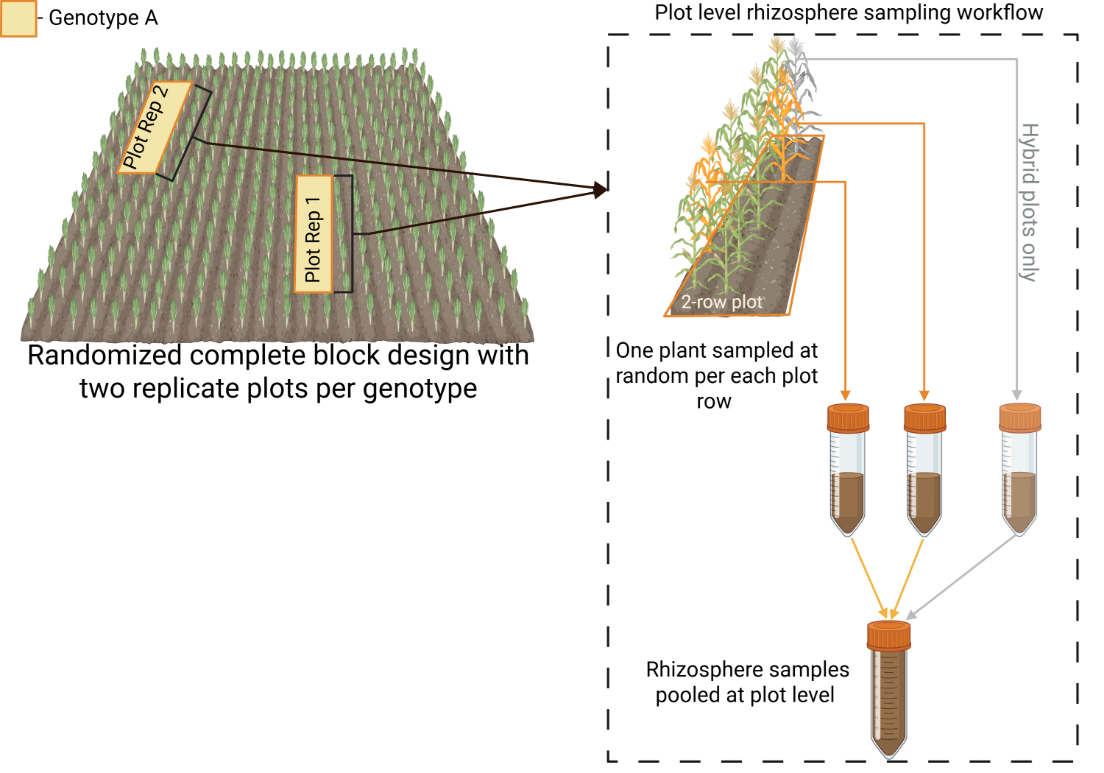

Note: To adequately characterize its rhizosphere community, each maize genotype was represented by two samples, with one collected from each of two replicate 2-row plots. For each replicate plot, one plant was randomly selected from each of the two rows, yielding a total of two rhizosphere samples, which were later pooled at the plot level. This sampling approach has previously been shown to both adequately capture field-scale spatial heterogeneity and introduce no statistically significant bias.^16^ Due to greater planting density, hybrid plots were sampled at three plants per plot, split between the two rows. The inbred lines were planted at a rate of 50 seeds per 2-row plot (25 seeds per row) with a row length of 5.2 m. The hybrid lines were planted at a rate of 80 seeds per 2-row plot (40 seeds per row) with an initial row length of 6.4 m, which was later reduced to 5.3 m. All plots had an inter-row spacing of 0.76 m.

Table S2 Sorting efficiency improvement based on the sorting reports

| Treatment | Sample Type | Sorting Efficiency (n=8) |
| --- | --- | --- |
| No Trypsin + EDTA | Sludge | 35.75% $\pm$4.2535 |
| Trypsin + 0.4mM EDTA | Soil | 85.5% $\pm$4.1833 |

Table S3 Potential polyphosphate (polyP)-accumulating organisms (PAOs) detected through TriFlow-Seq in EBPR sludge.

| Genus Name | Family/Order name | Genus within | polyP | PHA | Reference |
| --- | --- | --- | --- | --- | --- |
| *Tetrasphaera* | F_Intrasporangiaceae |  | o | o | ^17^ |
| *Dechloromonas* | F_Rhodocyclaceae |  | o | o | ^18^ |
| *Thauera* | F_Rhodocyclaceae |  |  |  | ^19^ |
| *Candidatus* Microthrix | F_ Microtrichaceae |  | o | x | ^20^ |
| *Ralstonia* | F_Burkholderiaceae |  | o | o | ^21^ |
| *Nitrospira* | F_Nitrospiraceae |  | o | x | ^22^ |
| *Alcaligenes* | F_Alcaligenaceae |  | o | o | ^23^ |
| *Novosphingobium* | F_ Sphingomonadaceae |  | o | o | ^24,25^ |
| *Zoogloea* | F_ Rhodocyclaceae |  | o | o | ^26^ |
| *Lautropia* | F_Burkholderiaceae |  | x | x | https://biocyc.org/GCF_900637555/organism-summary |
| *Planctomyces* | F_ Planctomycetaceae |  | x |  | https://www.ncbi.nlm.nih.gov/datasets/gene/GCF_001610835.1/?search=ppk |
| *Pirellula* | F_Pirellulaceae |  | x | x | https://biocyc.org/GCF_000025185/organism-summary |
| *Methylotenera* | F_Methylophilaceae |  | x | x | https://biocyc.org/GCF_000023705/organism-summary |
| *Thermomonas* | F_ Xanthomonadaceae |  | x | o | https://biocyc.org/GCF_014652775/organism-summary |
| *Ca*. Competibacter | F_Competibacteraceae |  | x | o | ^27^ |
| *Prosthecobacter* |  |  | x | o | https://biocyc.org/GCF_900167535/organism-summary |
| *Terrimonas* |  |  | x | x | https://biocyc.org/GCF_000425585/organism-summary |
| *Sorangium* | F_Polyangiaceae |  | x |  | https://biocyc.org/GCF_004135735/organism-summary |
| *Flavobacterium* | F_Flavobacteriaceae |  | x |  | https://www.ncbi.nlm.nih.gov/datasets/gene/GCF_000016645.1/?search=ppk |
| *Ferruginibacter* | F_ Chitinophagaceae |  | x |  | https://www.ncbi.nlm.nih.gov/datasets/gene/GCF_020042285.1/?search=ppk |
| *Dokdonella* | F_ Rhodanobacteraceae |  | x | x | <https://www.ncbi.nlm.nih.gov/datasets/gene/GCF_004342425.1/?search=phac>  <https://www.ncbi.nlm.nih.gov/datasets/gene/GCF_004342425.1/?search=ppk> |
| *Sulfuritalea* | F_ Rhodocyclaceae |  | x |  | https://www.ncbi.nlm.nih.gov/datasets/gene/GCF_000828635.1/?search=ppx |
| *Turneriella* | F_ Leptospiraceae |  | x |  | https://www.ncbi.nlm.nih.gov/datasets/gene/GCF_000266885.1/?search=ppk |
| *Haliangium* | F_ Haliangiaceae |  | x |  | https://www.ncbi.nlm.nih.gov/datasets/gene/GCF_000024805.1/?search=ppk |
| *Denitratisoma* | F_ Rhodocyclaceae |  | x | x | <https://www.ncbi.nlm.nih.gov/datasets/gene/GCF_902813185.1/?search=phac>  https://www.ncbi.nlm.nih.gov/datasets/gene/GCF_902813185.1/?search=ppx |
|  | O_Rhizobiales | Lichenibacterium | o | o | ^28^ |
|  | F_Alcaligenaceae | Alcaligenes | o | o | ^23^ |
|  | F_Comamonadaceae | Lampropedia; | o | o | ^18^ |
|  | F_Anaerolineaceae | Anaerolinea | x |  | https://www.ncbi.nlm.nih.gov/datasets/gene/GCF_001050195.2/?search=ppk |
|  | F_ Myxococcales | Myxococcus xanthus | o |  | ^29^ |
|  | F_ Saprospiraceae | OLB8 | x | x | ^30^ |
|  | F_ Hydrogenophilaceae | Hydrogenophilus thermoluteolus | x | o | ^31^ |
| Unclassified | F_Rhodocyclaceae | Dechloromonas | o | o | ^18^ |
|  | Obscuribacterales | Candidatus Obscuribacter | x | o | https://www.ncbi.nlm.nih.gov/datasets/gene/GCA_016712465.1/?search=ppk |
|  | F_Rhodobacteraceae | Rhodovulum | x | o | ^32^ |
|  | C_Planctomyces | *Gemmata* | x | x | ^33^ |
|  | O_Sphingobacteriales | *Sphingobacterium* | x |  | https://www.ncbi.nlm.nih.gov/datasets/gene/GCF_901482695.1/?search=ppk |
|  | F_Blastocatellaceae | Aridibacter | x |  | https://www.ncbi.nlm.nih.gov/datasets/gene/GCA_035513035.1/?search=ppk |
|  | F_Caldilineaceae | Caldilinea, Litorilinea | x |  | https://www.ncbi.nlm.nih.gov/datasets/gene/GCF_006569185.2/?search=ppk |
| Note: ‘x’ indicates gene potential for polyP and PHA accumulation according to reference genome; ‘o’ indicates experimental evidence of polyP and PHA accumulation through staining that reported in literature.  The color codes are the same with Figure 2. Red indicates the genus has been experimentally confirmed with PHA/polyP and PAO activities; Yellow indicates the presence of polyP and PHA; green indicates the the reference genome contains necessary genes for polyP and/or carbon storage synthesis; blue indicates some genera within this family have polyP accumulation ability | | | | | |

Table S4 Potential polyphosphate (polyP)-accumulating organisms (PAOs) detected through TriFlow-Seq in rhizosphere soil.

| Genus Name | Family/Order name | polyP | PHA | Reference |
| --- | --- | --- | --- | --- |
| *Halomonas* | Halomonadaceae | o | o | ^34^ |
| *Pseudomonas* | Pseudomonadaceae | o | o | ^35^ |
| *Steroidobacter* | Steroidobacteraceae | o | o | ^36^ |
| *Bacillus* | Bacillaceae |  |  | ^37^ |
| *Bradyrhizobium* | Nitrobacteraceae | o | o | ^36^ |
| *Hyphomicrobium* | Hyphomicrobiaceae | o | o | ^38^ |
|  | Vicinamibacteraceae | o | o | ^39^ |
| *Nitrospira* japonica | Nitrospiraceae | o | o | ^40^ |
| *Gaiella* | Gaiellaceae | x |  | https://www.ncbi.nlm.nih.gov/datasets/gene/GCF_003351045.1/?search=ppk |
| *Microvirga* | Methylobacteriaceae | x | x | <https://www.ncbi.nlm.nih.gov/datasets/gene/GCF_002741015.1/?search=ppk> https://www.ncbi.nlm.nih.gov/datasets/gene/GCF_002741015.1/?search=phac |
| *Nannocystis* | Nannocystaceae | x |  | https://www.ncbi.nlm.nih.gov/datasets/gene/GCF_016714285.1/?search=ppk |
|  | Nitrososphaeraceae |  |  |  |
| *Paenisporosarcina* |  | x | x | https://www.ncbi.nlm.nih.gov/datasets/gene/GCF_004367585.1/?search=phac |
|  | Planococcaceae | x |  |  |
| *Pseudarthrobacter* | Micrococcaceae | x |  | https://www.ncbi.nlm.nih.gov/datasets/gene/GCF_000022025.1/?search=ppk |
| *Ramlibacter* | Comamonadaceae | x | x | https://www.ncbi.nlm.nih.gov/datasets/gene/GCF_000215705.1/?search=ppk |
| *Skermanella aerolata* | Azospirillaceae | x | x | https://www.ncbi.nlm.nih.gov/datasets/gene/GCF_000936425.1/?search=ppk |
| *Ca*. Nitrocosmicus | Nitrososphaeraceae | x |  | https://www.ncbi.nlm.nih.gov/datasets/gene/GCF_000802205.1/?search=ppk |
| *Ca*. Udaeobacter | Chthoniobacteraceae | x |  | https://www.ncbi.nlm.nih.gov/datasets/gene/GCA_036499375.1/?search=ppk |
| Note: x indicates gene potential for polyP and PHA accumulation according to reference genome; ‘o’ indicates experimental evidence of polyP and PHA accumulation through staining that reported in literature.  Color codes are the same with Figure 2. Red indicates the genus has been experimentally confirmed with PHA/polyP and PAO activities; Yellow indicates the presence of polyP and PHA; Green indicates the the reference genome contains necessary genes for polyP and carbon storage synthesis; Blue indicates some genus within this Genes/Family have polyP accumulation ability | | | | |

References

(1)	Gu, A. Z.; Saunders, A.; Neethling, J. B.; Stensel, H. D.; Blackall, L. L. Functionally Relevant Microorganisms to Enhanced Biological Phosphorus Removal Performance at Full-Scale Wastewater Treatment Plants in the United States. *Water Environment Research* **2008**, *80* (8), 688–698. https://doi.org/10/b4n4ss.

(2)	Li, C.; Zeng, W.; Li, N.; Guo, Y.; Peng, Y.; Chao Li, a Wei Zeng, a Ning Li, a Yu Guo, a Y. P. Population Structure and Morphotype Analysis of “Candidatus Accumulibacter” Using Fluorescence In Situ HybridizationStaining-Flow Cytometry. *Applied and Environmental Microbiology* **2019**, *85* (February), 1–13.

(3)	Mori, F.; Nishimura, T.; Wakamatsu, T.; Terada, T.; Morono, Y. Simple In-Liquid Staining of Microbial Cells for Flow Cytometry Quantification of the Microbial Population in Marine Subseafloor Sediments. *Microbes and Environments* **2021**, *36* (3), 2–7. https://doi.org/10.1264/jsme2.ME21031.

(4)	Bartram, A. K.; Lynch, M. D. J.; Stearns, J. C.; Moreno-Hagelsieb, G.; Neufeld, J. D. Generation of Multi-Million 16S rRNA Gene Libraries from Complex Microbial Communities by Assembling Paired-End Illumina Reads. *Applied and environmental microbiology* **2011**, *77* (11), 3846–3852.

(5)	Quast, C.; Pruesse, E.; Yilmaz, P.; Gerken, J.; Schweer, T.; Yarza, P.; Peplies, J.; Glöckner, F. O. The SILVA Ribosomal RNA Gene Database Project: Improved Data Processing and Web-Based Tools. *Nucleic acids research* **2012**, *41* (D1), D590–D596.

(6)	Callahan, B. J.; McMurdie, P. J.; Holmes, S. P. Exact Sequence Variants Should Replace Operational Taxonomic Units in Marker-Gene Data Analysis. *ISME J* **2017**, *11* (12), 2639–2643. https://doi.org/10.1038/ismej.2017.119.

(7)	Callahan, B. J.; McMurdie, P. J.; Rosen, M. J.; Han, A. W.; Johnson, A. J. A.; Holmes, S. P. DADA2: High-Resolution Sample Inference from Illumina Amplicon Data. *Nat Methods* **2016**, *13* (7), 581–583. https://doi.org/10.1038/nmeth.3869.

(8)	Chao, A.; Gotelli, N. J.; Hsieh, T. C.; Sander, E. L.; Ma, K. H.; Colwell, R. K.; Ellison, A. M. Rarefaction and Extrapolation with Hill Numbers: A Framework for Sampling and Estimation in Species Diversity Studies. *Ecological Monographs* **2014**, *84* (1), 45–67. https://doi.org/10.1890/13-0133.1.

(9)	Li, H.-Z.; Bi, Q.; Yang, K.; Zheng, B.-X.; Pu, Q.; Cui, L. D2O-Isotope-Labeling Approach to Probing Phosphate-Solubilizing Bacteria in Complex Soil Communities by Single-Cell Raman Spectroscopy. *Anal. Chem.* **2019**, *91* (3), 2239–2246. https://doi.org/10.1021/acs.analchem.8b04820.

(10)	Li, G.; Wu, C.; Wang, D.; Srinivasan, V.; Dy, J. G.; Gu, A. Z. Machine Learning-Based Determination of Sampling Depth for Complex Environmental Systems: Case Study with Single-Cell Raman Spectroscopy Data in EBPR Systems. *Environ. Sci. Technol.* **2022**, *56* (18), 13473–13484. https://doi.org/10.1021/acs.est.1c08768.

(11)	Li, Y.; Cope, H. A.; Rahman, S. M.; Li, G.; Nielsen, P. H.; Elfick, A.; Gu, A. Z. Toward Better Understanding of EBPR Systems via Linking Raman-Based Phenotypic Profiling with Phylogenetic Diversity. *Environmental Science and Technology* **2018**, *52* (15), 8596–8606. https://doi.org/10.1021/acs.est.8b01388.

(12)	Wang, D.; Li, Y.; Cope, H. A.; Li, X.; He, P.; Liu, C.; Li, G.; Rahman, S. M.; Tooker, N. B.; Bott, C. B.; Onnis-Hayden, A.; Singh, J.; Elfick, A.; Marques, R.; Jessen, H. J.; Oehmen, A.; Gu, A. Z. Intracellular Polyphosphate Length Characterization in Polyphosphate Accumulating Microorganisms (PAOs): Implications in PAO Phenotypic Diversity and Enhanced Biological Phosphorus Removal Performance. *Water Research* **2021**, *206*, 117726. https://doi.org/10.1016/j.watres.2021.117726.

(13)	Majed, N.; Matthäus, C.; Diem, M.; Gu, A. Z. Evaluation of Intracellular Polyphosphate Dynamics in Enhanced Biological Phosphorus Removal Process Using Raman Microscopy. *Environmental science & technology* **2009**, *43* (14), 5436–5442.

(14)	Majed, N.; Chernenko, T.; Diem, M.; Gu, A. Z. Identification of Functionally Relevant Populations in Enhanced Biological Phosphorus Removal Processes Based on Intracellular Polymers Profiles and Insights into the Metabolic Diversity and Heterogeneity. *Environmental science & technology* **2012**, *46* (9), 5010–5017.

(15)	Majed, N.; Gu, A. Z. Application of Raman Microscopy for Simultaneous and Quantitative Evaluation of Multiple Intracellular Polymers Dynamics Functionally Relevant to Enhanced Biological Phosphorus Removal Processes. *Environmental science & technology* **2010**, *44* (22), 8601–8608.

(16)	Allen, W. J.; Sapsford, S. J.; Dickie, I. A. Soil Sample Pooling Generates No Consistent Inference Bias: A Meta‐analysis of 71 Plant–Soil Feedback Experiments. *New Phytologist* **2021**, *231* (4), 1308–1315.

(17)	Kristiansen, R.; Nguyen, H. T. T.; Saunders, A. M.; Nielsen, J. L.; Wimmer, R.; Le, V. Q.; McIlroy, S. J.; Petrovski, S.; Seviour, R. J.; Calteau, A.; Nielsen, K. L.; Nielsen, P. H. A Metabolic Model for Members of the Genus Tetrasphaera Involved in Enhanced Biological Phosphorus Removal. *ISME Journal* **2013**, *7* (3), 543–554. https://doi.org/10.1038/ismej.2012.136.

(18)	Ruiz-Haddad, L.; Ali, M.; Pronk, M.; van Loosdrecht, M. C. M.; Saikaly, P. E. Demystifying Polyphosphate-Accumulating Organisms Relevant to Wastewater Treatment: A Review of Their Phylogeny, Metabolism, and Detection. *Environmental Science and Ecotechnology* **2024**, *21*, 100387. https://doi.org/10.1016/j.ese.2024.100387.

(19)	Ren, T.; Chi, Y.; Wang, Y.; Shi, X.; Jin, X.; Jin, P. Diversified Metabolism Makes Novel *Thauera* Strain Highly Competitive in Low Carbon Wastewater Treatment. *Water Research* **2021**, *206*, 117742. https://doi.org/10.1016/j.watres.2021.117742.

(20)	Nierychlo, M.; Singleton, C. M.; Petriglieri, F.; Thomsen, L.; Petersen, J. F.; Peces, M.; Kondrotaite, Z.; Dueholm, M. S.; Nielsen, P. H. Low Global Diversity of Candidatus Microthrix, a Troublesome Filamentous Organism in Full-Scale WWTPs. *Front Microbiol* **2021**, *12*, 690251. https://doi.org/10/gstz8q.

(21)	Tumlirsch, T.; Sznajder, A.; Jendrossek, D. Formation of Polyphosphate by Polyphosphate Kinases and Its Relationship to Poly(3-Hydroxybutyrate) Accumulation in Ralstonia Eutropha Strain H16. *Applied and Environmental Microbiology* **2015**, *81* (24), 8277–8293. https://doi.org/10.1128/AEM.02279-15.

(22)	Lebedeva, E. V.; Alawi, M.; Maixner, F.; Jozsa, P.-G.; Daims, H.; Spieck, E. Physiological and Phylogenetic Characterization of a Novel Lithoautotrophic Nitrite-Oxidizing Bacterium, ‘Candidatus Nitrospira Bockiana.’ *International Journal of Systematic and Evolutionary Microbiology* **2008**, *58* (1), 242–250. https://doi.org/10.1099/ijs.0.65379-0.

(23)	Doi, Y.; Kawaguchi, Y.; Nakamura, Y.; Kunioka, M. Nuclear Magnetic Resonance Studies of Poly(3-Hydroxybutyrate) and Polyphosphate Metabolism in Alcaligenes Eutrophus. *Applied and Environmental Microbiology* **1989**, *55* (11), 2932–2938. https://doi.org/10.1128/aem.55.11.2932-2938.1989.

(24)	Smit, A.-M.; Strabala, T. J.; Peng, L.; Rawson, P.; Lloyd-Jones, G.; Jordan, T. W. Proteomic Phenotyping of Novosphingobium Nitrogenifigens Reveals a Robust Capacity for Simultaneous Nitrogen Fixation, Polyhydroxyalkanoate Production, and Resistance to Reactive Oxygen Species. *Applied and Environmental Microbiology* **2012**, *78* (14), 4802–4815. https://doi.org/10.1128/AEM.00274-12.

(25)	Belmok, A.; de Almeida, F. M.; Rocha, R. T.; Vizzotto, C. S.; Tótola, M. R.; Ramada, M. H. S.; Krüger, R. H.; Kyaw, C. M.; Pappas, G. J. Genomic and Physiological Characterization of Novosphingobium Terrae Sp. Nov., an Alphaproteobacterium Isolated from Cerrado Soil Containing a Mega-Sized Chromid. *Braz J Microbiol* **2023**, *54* (1), 239–258. https://doi.org/10.1007/s42770-022-00900-4.

(26)	Oshiki, M.; Satoh, H.; Mino, T.; Onuki, M. PHA-Accumulating Microorganisms in Full-Scale Wastewater Treatment Plants. *Water Science and Technology* **2008**, *58* (1), 13–20. https://doi.org/10/frbtcv.

(27)	McIlroy, S. J.; Albertsen, M.; Andresen, E. K.; Saunders, A. M.; Kristiansen, R.; Stokholm-Bjerregaard, M.; Nielsen, K. L.; Nielsen, P. H. ‘Candidatus Competibacter’-Lineage Genomes Retrieved from Metagenomes Reveal Functional Metabolic Diversity. *ISME J* **2014**, *8* (3), 613–624. https://doi.org/10.1038/ismej.2013.162.

(28)	Pankratov, T. A.; Grouzdev, D. S.; Patutina, E. O.; Kolganova, T. V.; Suzina, N. E.; Berestovskaya, J. J. Lichenibacterium Ramalinae Gen. Nov, Sp. Nov., Lichenibacterium Minor Sp. Nov., the First Endophytic, Beta-Carotene Producing Bacterial Representatives from Lichen Thalli and the Proposal of the New Family Lichenibacteriaceae within the Order Rhizobiales. *Antonie van Leeuwenhoek* **2020**, *113* (4), 477–489. https://doi.org/10.1007/s10482-019-01357-6.

(29)	Kurashita, H.; Hatamoto, M.; Tomita, S.; Yamaguchi, T.; Narihiro, T.; Kuroda, K. Comprehensive Insights into Potential Metabolic Functions of Myxococcota in Activated Sludge Systems. *Microbes and Environments* **2024**, *39* (4). https://doi.org/10.1264/jsme2.ME24068.

(30)	Kondrotaite, Z.; Valk, L. C.; Petriglieri, F.; Singleton, C.; Nierychlo, M.; Dueholm, M. K. D.; Nielsen, P. H. Diversity and Ecophysiology of the Genus OLB8 and Other Abundant Uncultured Saprospiraceae Genera in Global Wastewater Treatment Systems. *Front Microbiol* **2022**, *13*, 917553. https://doi.org/10.3389/fmicb.2022.917553.

(31)	Nguyen, T. H.; Ishizuna, F.; Sato, Y.; Arai, H.; Ishii, M. Physiological Characterization of Poly-β-Hydroxybutyrate Accumulation in the Moderately Thermophilic Hydrogen-Oxidizing Bacterium Hydrogenophilus Thermoluteolus TH-1. *J Biosci Bioeng* **2019**, *127* (6), 686–689. https://doi.org/10.1016/j.jbiosc.2018.11.011.

(32)	Higuchi-Takeuchi, M.; Motoda, Y.; Kigawa, T.; Numata, K. Class I Polyhydroxyalkanoate Synthase from the Purple Photosynthetic Bacterium Rhodovulum Sulfidophilum Predominantly Exists as a Functional Dimer in the Absence of a Substrate. *ACS Omega* **2017**, *2* (8), 5071–5078. https://doi.org/10.1021/acsomega.7b00667.

(33)	Boedeker, C.; Schüler, M.; Reintjes, G.; Jeske, O.; van Teeseling, M. C. F.; Jogler, M.; Rast, P.; Borchert, D.; Devos, D. P.; Kucklick, M.; Schaffer, M.; Kolter, R.; van Niftrik, L.; Engelmann, S.; Amann, R.; Rohde, M.; Engelhardt, H.; Jogler, C. Determining the Bacterial Cell Biology of Planctomycetes. *Nat Commun* **2017**, *8* (1), 14853. https://doi.org/10.1038/ncomms14853.

(34)	Nguyen, H. T. T.; Nielsen, J. L.; Nielsen, P. H. ‘Candidatus Halomonas Phosphatis’, a Novel Polyphosphate-Accumulating Organism in Full-Scale Enhanced Biological Phosphorus Removal Plants. *Environmental Microbiology* **2012**, *14* (10), 2826–2837. https://doi.org/10.1111/j.1462-2920.2012.02826.x.

(35)	Tobin, K. M.; McGrath, J. W.; Mullan, A.; Quinn, J. P.; O’Connor, K. E. Polyphosphate Accumulation by Pseudomonas Putida CA-3 and Other Medium-Chain-Length Polyhydroxyalkanoate-Accumulating Bacteria under Aerobic Growth Conditions. *Applied and Environmental Microbiology* **2007**, *73* (4), 1383–1387. https://doi.org/10.1128/AEM.02007-06.

(36)	Sharma, V.; Siedenburg, G.; Birke, J.; Mobeen, F.; Jendrossek, D.; Prakash, T. Metabolic and Taxonomic Insights into the Gram-Negative Natural Rubber Degrading Bacterium Steroidobacter Cummioxidans Sp. Nov., Strain 35Y. *PLOS ONE* **2018**, *13* (5), e0197448. https://doi.org/10.1371/journal.pone.0197448.

(37)	Acosta-Cortés, A. G.; Martinez-Ledezma, C.; López-Chuken, U. J.; Kaushik, G.; Nimesh, S.; Villarreal-Chiu, J. F. Polyphosphate Recovery by a Native Bacillus Cereus Strain as a Direct Effect of Glyphosate Uptake. *The ISME Journal* **2019**, *13* (6), 1497–1505. https://doi.org/10.1038/s41396-019-0366-3.

(38)	Martinez, R. J.; Wu, C. H.; Beazley, M. J.; Andersen, G. L.; Conrad, M. E.; Hazen, T. C.; Taillefert, M.; Sobecky, P. A. Microbial Community Responses to Organophosphate Substrate Additions in Contaminated Subsurface Sediments. *PLOS ONE* **2014**, *9* (6), e100383. https://doi.org/10.1371/journal.pone.0100383.

(39)	Kristensen, J. M.; Singleton, C.; Clegg, L.-A.; Petriglieri, F.; Nielsen, P. H. High Diversity and Functional Potential of Undescribed “Acidobacteriota” in Danish Wastewater Treatment Plants. *Frontiers in Microbiology* **2021**, *12*, 643950. https://doi.org/10.3389/fmicb.2021.643950.

(40)	Yang, Y.; Daims, H.; Liu, Y.; Herbold, C. W.; Pjevac, P.; Lin, J.-G.; Li, M.; Gu, J.-D. Activity and Metabolic Versatility of Complete Ammonia Oxidizers in Full-Scale Wastewater Treatment Systems. *mBio* **2020**, *11* (2), e03175-19. https://doi.org/10/gn5874.
